# Diversity of motility patterns in benthic diatoms

**DOI:** 10.1101/2024.12.23.630140

**Authors:** Karen Grace Bondoc-Naumovitz, Emanuele Crosato, Kirsty Y. Wan

## Abstract

Diatoms, a highly successful group of photosynthetic algae, contribute to a quarter of global primary production. Many species are motile, despite having no appendages and a completely rigid cell body. Cells move to seek out nutrients, locate mating partners, and undergo vertical migration. To explore the natural diversity of diatom motility, we perform a comparative study across five common biofilm-forming species. Combining morphological measurements with high-resolution cell tracking, we establish how gliding movements relate to the morphology of the raphe – a specialised slit in the cell wall responsible for motility generation. Our detailed analyses reveal that cells exhibit a rich but species-dependent phenotype, switching stochastically between four stereotyped motility states. We model this behaviour and use stochastic simulations to predict how heterogeneity in microscale navigation patterns leads to differences in long-time diffusivity and dispersal. In a representative species, we extend these findings to quantify diatom gliding in complex, naturalistic 3D environments, suggesting that cells may exploit these distinct motility signatures to achieve niche segregation in nature.

---

From the smallest bacterium to the largest animals, locomotion is a conserved trait that allows organisms to thrive, and diversify in new habitats [1, 2, 3]. Microscopic organisms display an overwhelming array of movement mechanisms for navigating in 2D and 3D, yet, the same mechanism can also be used by different organisms due to convergent evolution [4]. Gliding motility, where cells move actively while adhering to substrates, is present in diverse phylogenetically unrelated species, including myxobacteria, mollicutes, cyanobacteria, apicomplexans [4, 5, 6, 7]. Many benthic diatoms modulate their gliding motility in response to environmental changes [8, 9, 10, 11, 12, 13, 14, 15], yet the cells have no appendages and cannot change shape due to their completely rigid silica cell walls. This unusual motility is in contrast to other motile organisms that use specialised appendages such as cilia, flagella and pili [16, 17] or temporary protrusions in the case of amoeboid movement [18].

Diatom gliding represents a unique and mechanistically interesting system for understanding cell migration on surfaces without shape changes. Gliding involves the secretion of mucus-like extracellular polymeric substances (EPS) through a raphe – a specialised slit in the cell wall (Fig. 1a), though the exact molecular mechanism remains disputed [19, 20]. Movement is likely driven by an actomyosin system [21, 22, 23, 24], where myosin translocation coincides with cell movement in the opposite direction (Fig. 1b). In contrast to gliding in most other eukaryotic cells where actin filaments are remodelled [25, 26, 27], here actin bundles are fixed (running parallel to the raphe), again emphasizing a possible functional role of the raphe.

**Figure 1.**
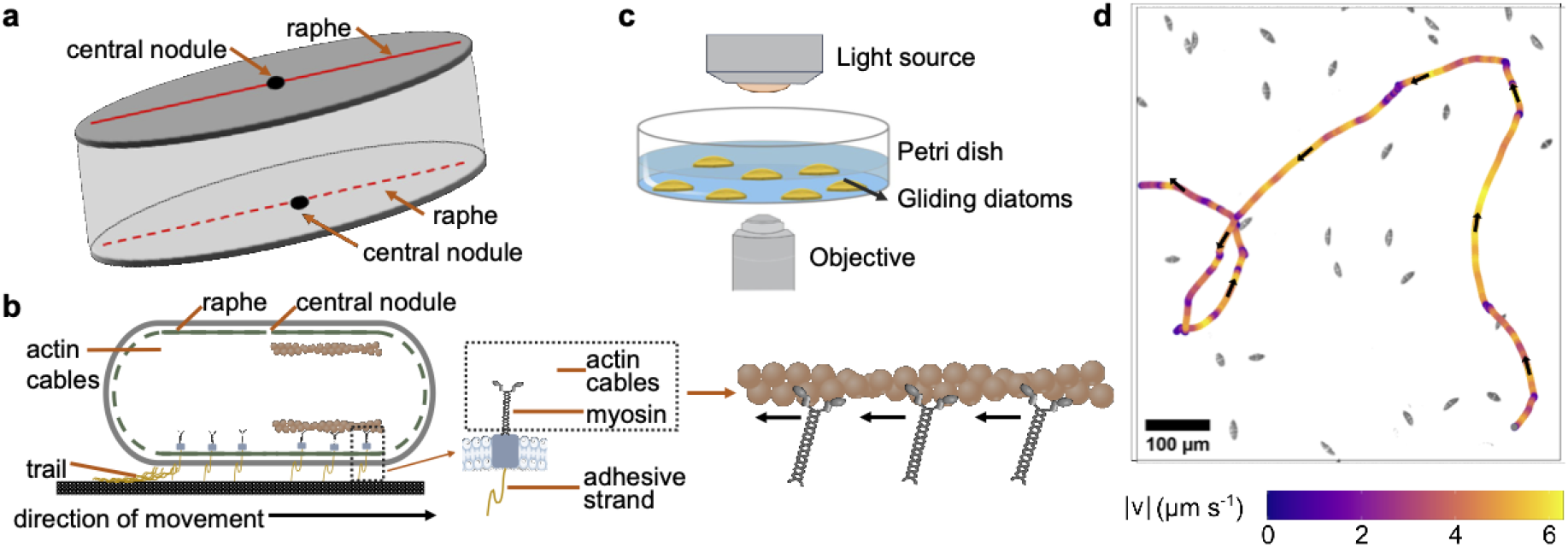
Diatom morphology and gliding motility assay. **a** Schematic of a diatom. The raphe (red solid and dotted lines), a specialised slit in the cell wall implicated in cell movement, runs along the whole length of the valve either continuously or interrupted by the central nodule. The raphe is present on both the top and bottom valves of the cell. **b** Hypothesised mechanism of diatom gliding. Secreted adhesive strands are connected via transmembrane components to the actin-myosin complex underlying the raphe. The cell moves in the opposite direction of the force supplied by myosin motors acting along the actin cables (modified image from Molino and Wetherbee [22]). **c** Experimental set-up. Diatoms cultured on a glass-bottomed dish (35 mm) were imaged using an inverted light microscope. **d** A sample gliding trajectory of the diatom *Seminavis robusta*, colour-coded by instantaneous glide speed (arrows show the direction of motion).

As a major constituent of benthic ecosystems, diatom communities play a key role in global carbon fixation and nutrient cycling [28, 29, 30]. They actively choose surfaces for attachment and gliding, and exploit fine-scale spatial and temporal heterogeneity in the environment such as light, dissolved nutrients, and mating pheromones [8, 9, 10, 14]. Some species also exhibit vertical diel migration through the photic zone (~500 *µ*m – 3 mm), an efficient process consuming only 0.03 pcal (or 0.0001% glucose consumption) for a cell moving upwards or downwards over a distance of 400 *µ*m [31, 32, 33]. Despite the significance of active motility in influencing species dispersal and ecosystem functioning, our knowledge of individual cell behaviours at the microscopic scale is limited [34, 35, 20, 21]. Previous studies have focused on resolving the migration and composition of diatom communities [28, 36, 37].

To address this knowledge gap, we present a comprehensive multispecies analysis of diatom motility, revealing an unprecedented diversity in motility patterns which may be attributed to differences in diatom body morphology and ecosystem habitat. Through quantitative single-cell imaging and tracking experiments (Figs. 1c,d, 3a, b), we show that diatoms exhibit a basic set of reproducible behaviours, or stereotyped states, but diverse gliding trajectories can be generated by changing the associated transition rates between states. This allows us to construct a generic model of diatom motility that effectively captures the detailed microscopic features of cell movement and predicts their long-time dispersal. To establish whether these features of diatom motility persist within complex, naturalistic environments, we also perform 3D motility tracking in a representative diatom - the first experiment of this kind. Our results emphasize the functional morphology of the raphe as a driver of divergent diatom gliding behaviours exhibited by distinct genera.

## Results

### Diatom raphe morphology influences gliding motility

We study five representative diatoms commonly found in benthic biofilms: *Cylindrotheca closterium* (hereafter *Cylindrotheca*), *Halamphora sp*. (hereafter *Halamphora*), *Nitzschia ovalis* (hereafter *Nitzschia*), *Pleurosigma sp*. (hereafter *Pleurosigma*), and *Seminavis robusta* (hereafter *Seminavis*). See Methods and SI Table 1 for details of the culturing and experimental procedures. The species were found to have distinctive body/raphe morphologies which we hypothesize to influence motility. Cells vary greatly in length from 5–85 *µ*m. We prepared samples for scanning electron microscopy (SEM) to visualise raphe shape and estimate raphe curvature *κ*_*r*_ by manual tracing (Fig. 2, SI Fig. 2; Methods).

**Figure 2.**
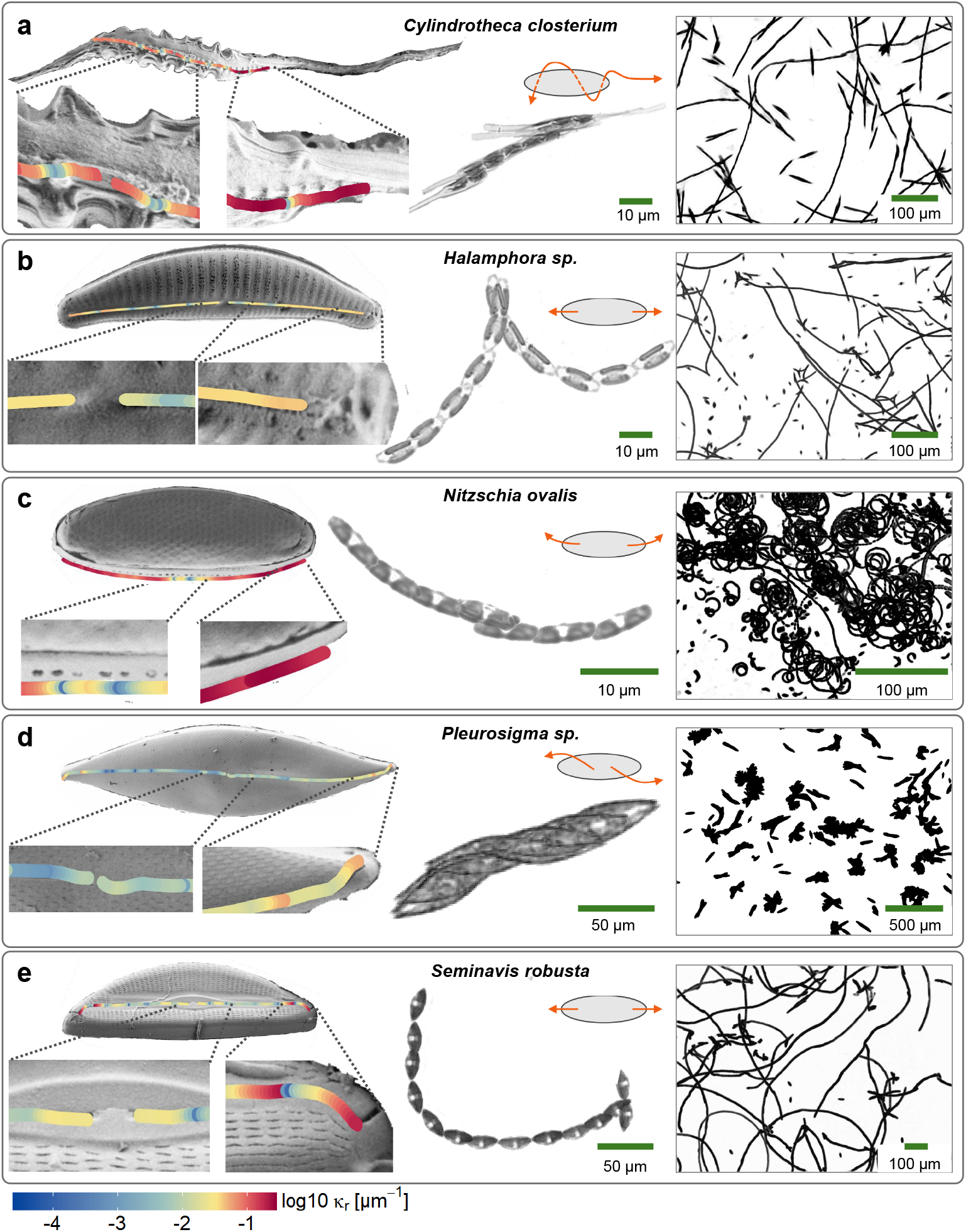
Raphe morphology and track patterns of five biofilm-forming diatoms. From left to right: scanning electron microscopy (SEM) of each diatom with raphe highlighted (colour overlay denotes the raphe curvature *κ*_*r*_), sample brightfield image of a moving cell (inset shows a schematic on the motion), and maximum intensity projection of a diatom population for **a** *Cylindrotheca closterium*, **b** *Halamphora sp*., **c** *Nitzschia ovalis*, **d** *Pleurosigma sp*., and **e** *Seminavis robusta*.

All five diatoms are biraphid pennates with two raphe systems on both valves, but they differ significantly in body shape and raphe placement [38] (Fig. 2). Biraphids may be symmetric (*Pleurosigma*) or asymmetric (*Seminavis, Halamphora*), depending on whether the valves are positioned symmetrically relative to the transpical axis of the diatom. More unusual raphe morphologies are found in *Cylindrotheca* and *Nitzschia*, both nitzschoid diatoms. *Cylindrotheca* has a 3D helical raphe spiralling along a needle-shaped cell body [35], which has higher curvature compared to the other diatoms (Fig. 1a). The raphe canal also has a non-smooth undulating surface. The raphe of *Nitzschia* lies inside a keel or canal with pores leading to the surface [38] (Fig. 1c). Both *Halamphora* and *Seminavis* have relatively straight raphes running from the central nodule to both ends (Figs. 2b,e). The distal raphe of *Seminavis* is slightly deflected to one side of the valve. In *Pleurosigma* the raphe follows the distinctive sigmoidal shape of the valves, with the distal raphe ends deflected towards opposite sides of the mantle (Fig. 2d).

The shape of the raphe, which is physically constrained by the cell body, is reflected in the trajectories (Fig. 2, columns 2 & 3). We explore this quantitatively by performing high-resolution singlecell tracking experiments (n=4121 across 5 species). Diatoms were cultured on glass-bottomed plastic microwells and allowed to establish an early biofilm to ensure surface conditions are primed for motility assays. Only a subset of the diatoms are motile (SI Fig. S1). Various gliding patterns were observed (SI Videos 1-5). Most cells followed straight or slightly curved tracks with frequent changes in direction (Fig. 2, second column). We measured the diatom’s centroid position **r**(*t*) = [*x*(*t*), *y*(*t*)] (spot tracking) and body orientation *θ*(*t*) (ellipse fitting of the segmented diatom image) simultaneously (Fig. 3a). From this, we calculated its translational velocity **v**(*t*), resolved into components *v*_∥_(*t*) and *v*_⊥_(*t*) parallel and perpendicular to the cell body, and rotational velocity *ω*(*t*) (Methods; Fig. 3b). Cell orientation angles were corrected for jump discontinuities.

**Figure 3.**
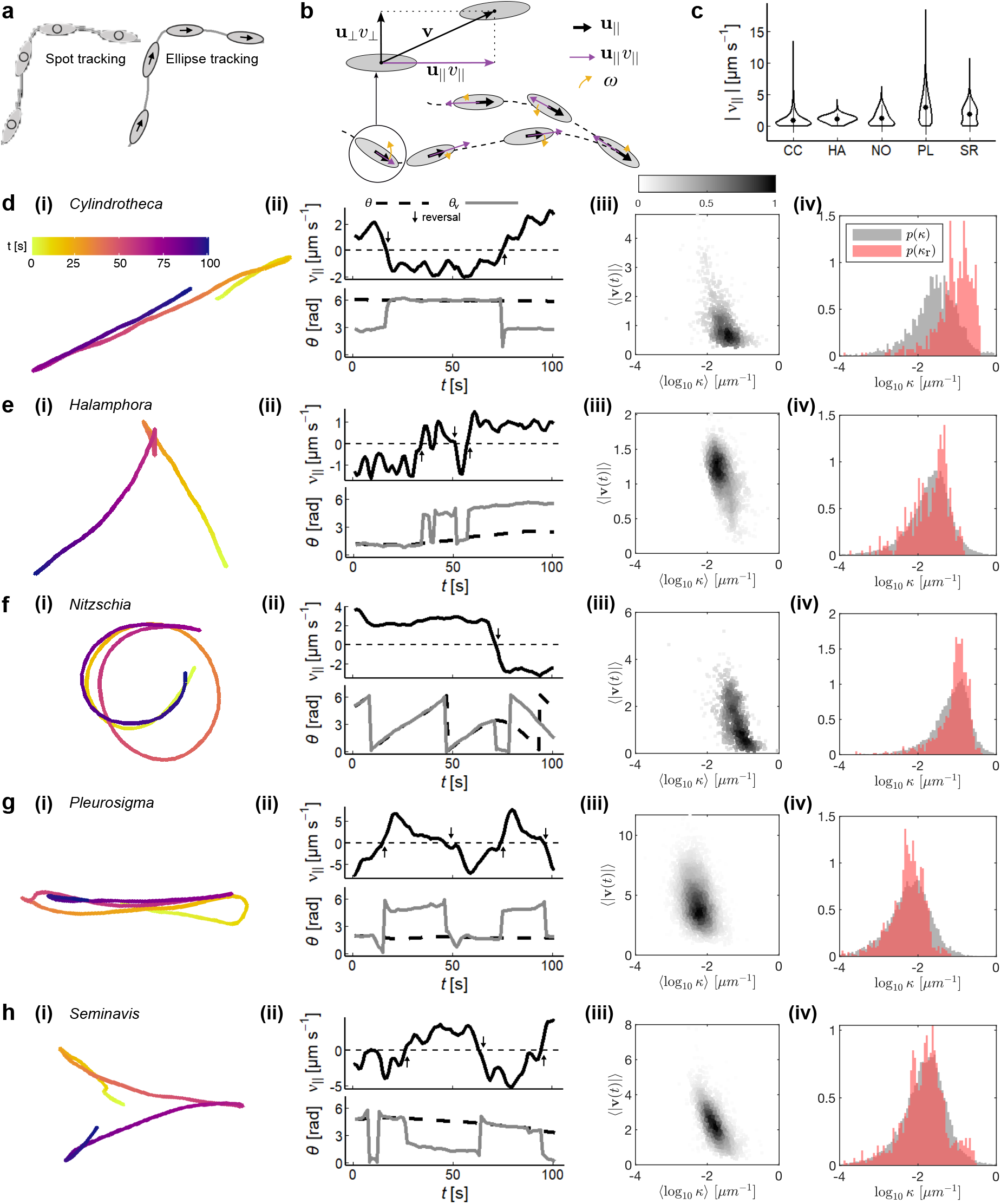
Gliding characteristics of diatoms. **a** Spot/centroid tracking and ellipse tracking were used to obtain the cell’s coordinate position, **r**(*t*) = [*x*(*t*), *y*(*t*)], and the orientation, *θ*, respectively. **b** From this, tangential velocity, *v*_∥_(*t*), and angular velocity, *ω*(*t*) were calculated. **c** Violin plot showing the distribution of tangential speed, |*v*_∥_|, of each diatom species. **d–h** Track characterisation. From left to right: (i) Representative trajectories coloured by time; (ii) Temporal dynamics of sample tracks highlighting the relationship between tangential velocity, *v*_∥_(*t*), cell orientation, *θ*(*t*), and direction of movement, *θ*_*v*_ (*t*); (iii) Bivariate histogram of the per-glide curvature ⟨log_10_ *κ*⟩ and ⟨|**v**|⟩; (iv) Histograms of track curvature *κ* and raphe *κ*_*r*_ for the ‘average’ diatom (see main text). The full distributions are shown in SI Fig. 5.

Diatom gliding is slow and jerky (< 20*µ*m *s*^−1^) compared to many other motile microbes (Fig. 3c). Representative trajectories are plotted in Fig. 3d-h:i, and colour-coded by time. *Halamphora* and *Seminavis* which have straight raphes predominantly form straight tracks with occasional turns and reverses. Meanwhile, *Cylindrotheca* glides with very frequent reversals (SI Videos 1, 2, 5). This intermittent behaviour may result from its twisted raphe since only a subsection can adhere to the surface at any given time. The largest diatom *Pleurosigma* does not reverse sharply but instead makes wider, hairpin-like reverses due to the sigmoidal raphe ends. Finally, *Nitzschia* forms either clockwise or counterclockwise circular trajectories. Comparing body orientation *θ*(*t*) with velocity direction *θ*_*v*_(*t*) reveals that straight glides, either forwards or backwards, coincide with near-perfect alignment of the diatom body with the track (Fig. 3d-h:*ii*). Instead, reversals are polarity switches where a gliding cell comes to rest (*v*(*t*) crosses zero) and immediately moves off in the opposite direction. These appear as ±*π* jumps in *θ*_*v*_(*t*) but the cell-fixed orientation *θ*(*t*) remains continuous.

Gliding, which corresponds to relatively straight paths, can be isolated from non-glides using a curvature threshold (SI Fig. 3). For each glide event, the mean curvature ⟨*κ*⟩ (see Methods and SI Fig. 4) and mean glide speed ⟨|*v*(*t*)|⟩ of individual glides are inversely related (Fig. 3d-h:*iii*), meaning that straight path segments coincides with fast movement, whereas higher curvature segments correspond to slower movement and transitions (see next section).

Next, we compare the histograms of *κ* (from track segments during glides) and *κ*_*r*_ (from SEM images) for each species (Fig. 3d-h:*iv*). We consider glide segments whose mean speed lies between the 45th and 55th percentiles (see Methods; SI Fig. 4), to select for diatoms *of average size*, which is most likely to be represented in the smaller SEM dataset. To see why average speed correlates with diatom length, we note that within the same species longer diatoms should have straighter raphes, and raphe curvature correlates negatively with speed (Fig. 3d-h:*iii*). Remarkably, the distributions match for all species, suggesting that diatoms glide on paths whose local curvature is determined by their raphes. For *Nitzschia*, which performs circular loops, the preferred glide curvature is most prominent. The agreement is less good for *Cylindrotheca*, partly because we cannot accurately trace 3D raphes from 2D projections in the SEM images. For this diatom, *κ*_*r*_ is generally higher than that of *κ*, likely because, only some parts of its helical raphe are aligned with the substrate.

Overall, these findings reveal that raphe morphology directly influences characteristics of the gliding paths of diatoms.

### Classification of diatom motility phenotypes

Despite major differences in trajectory morphology across the five species, we observed that the diatoms undergo just four stereotyped behaviours, namely Glide, Reverse, Pivot, and Stop (Fig. 4a). Discretizing motility patterns into low-dimensional gaits/states and studying transitions between these states allows us to quantify how behaviour changes over time, and make comparisons across species [5, 39, 40]. Glides are associated with persistent movement in a certain direction, involving motion along the body-axis direction (as discussed above). The circular motion of *Nitzschia* is also considered gliding in this interpretation. A reversal is a sudden event during which the diatom switches to gliding with its trailing end, which becomes the new leading end. Pivots occur when a diatom that is only partially adhered to a surface rotates about using the trailing end. Finally, a stop corresponds to cessation of movement.

**Figure 4.**
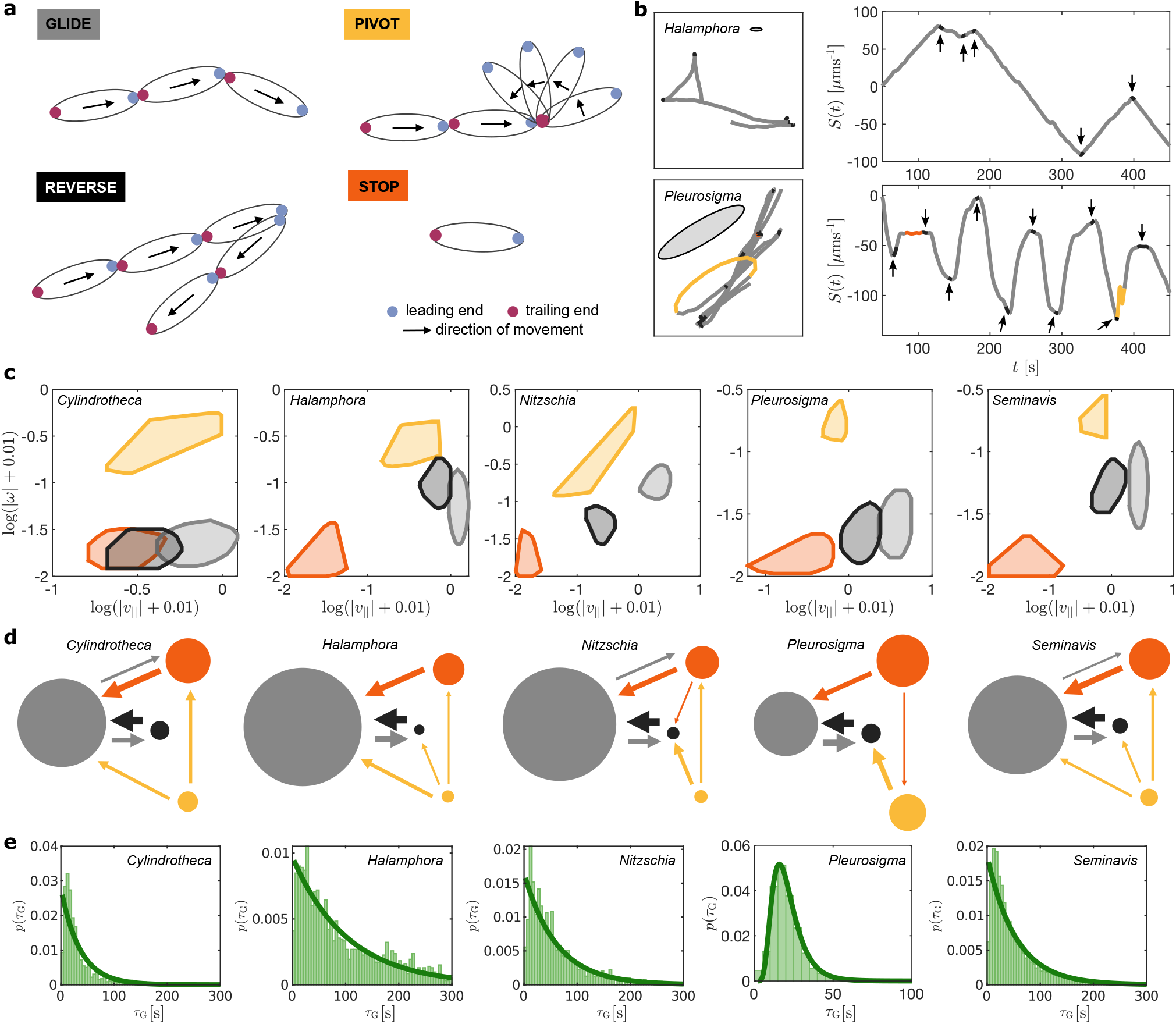
Four-state Markov model of diatom motility. **a** Definition of the four possible states. **b** Example of trajectory segmentation into motility states: the left panels – sample trajectories for *Halamphora* (top) and *Pleurosigma* (bottom), colour-coded based on the motility state (ellipses indicate the average size of each diatom), right panels – respective time evolution of *S*(*t*) (arrows highlight the reversals). **c** The four states visualised in the {|*v*_||_(*t*)|, |*ω*(*t*)|} parameter space. The region delineating each state is constructed from the bivariate histogram (logarithmic scale) (shifted by 0.01 for better visualisation of small values) after applying a smoothing filter and calculating the convex hull (97.5-percentile). **d** Distinct transition networks for each diatom species. Circles represent the states, arrows the transitions between states. The size of each circle scales with the mean dwell time per state, while the thickness of each arrow scales with the pair-wise transition probability between states (transitions with probability ≤ 0.2 have been pruned for ease of visualisation). **e** Distributions of gliding duration with fit to exponential distributions (*Cylindrotheca, Halamphora, Nitzschia* and *Seminavis*)) or lognormal distribution (*Pleurosigma*).

We classified the state of a diatom at every time frame (see Methods). The output is illustrated with two example tracks in Fig. 4b, where the trajectories are coloured by state, and plotted alongside the corresponding cumulative tangential velocity defined by 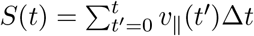. More example tracks can be found in SI Fig. 6. *Halamphora* glides at a very stable speed (*S*(*t*) has almost constant positive or negative slope). Glides are interspersed with random reversals (arrowheads), after which the preferred glide speed is quickly reestablished. Similar behaviour is observed for *Seminavis* and *Nitzschia* (although with chiral motion) and, to a certain extent, also for *Cylindrotheca*, whose movements are more irregular. *Pleurosigma* exhibits quasi-periodic interruptions to movement, accelerating/decelerating more gradually resulting in very constrained tracks.

Across all species, the four states are well differentiated by the diatoms’ tangential speed *v*_||_(*t*) and rotational speed *ω*(*t*). The four states occupy similar regions of the {|*v*_||_(*t*)|, |*ω*(*t*)|}-parameter space (Fig. 4c), despite differences in absolute values across species. A greater overlap is observed for *Cylindrotheca*, due to its low and erratic speed.

We used a four-state Markov chain to estimate the statistics of state transitions, where all non-self transitions are assumed possible except those leaving the Reverse state towards any other state than Glide, since a reversal implies the reestablishment of persistent, directed motion. We estimated mean dwell times and pairwise transition probabilities from the data and represent these as transition networks (Fig. 4d) (see Methods; SI Tables 2–11). All diatoms are found predominantly in the Glide state, often accounting for 55–75% of the time, the rest is spent in the Stop, Pivot, and Reverse states, in descending order (SI Fig. 7). For all diatoms, transitions between Glide and Reverse, or between Glide and Stop, are highly likely. The characteristic transition probabilities and mean dwelling times vary greatly across species, especially for Glide. While Glide durations are strongly exponentially distributed for most diatoms (Fig. 4e), *Pleurosigma* exhibits frequent and periodic reversals which we fit instead to a lognormal distribution. *Pleurosigma* is the only species showing oscillatory behaviour, with a periodicity of approximately 23 s (SI Table 10).

We showed that different diatoms exhibit distinct but stereotyped gliding behaviours that are fully captured by stochastic transitions between just four underlying states. These states may be a conserved feature of all raphe-based motility systems, while the main difference between species is that they can access distinct motility patterns by uniquely combining these gaits according to different transition rates.

### Stochastic model of diatom gliding

Next, we develop simulations of diatom gliding motility, with physical parameters calibrated directly using trajectories from each species. The stochasticity of diatom movement is captured using Stochastic Differential Equations (SDEs) for the evolution of the 2D position **r**(*t*) = [*x*(*t*), *y*(*t*)] and the orientation *θ*(*t*): d**r**(t) = f(*θ*(*t*), *s*(*t*), *t*)d*t* + **u**_||_(*t*)*v*_||*σ*_ d*W*_tra_(*t*), d*θ*(*t*) = *g*(*s*(*t*), *t*)d*t* + *ω*_*σ*_d*W*_rot_(*t*), where **u**_||_ = [cos *θ*(*t*), sin *θ*(*t*)], *W*_tra_ and *W*_rot_ are two uncorrelated Wiener processes so that *E*[d*W*_*i*_(*t*)d*W*_*j*_(*t*^*′*^)] = *δ*_*ij*_*δ*(*t* − *t*^*′*^), where *i* and *j* are indices for the labels ‘tra’ and ‘rot’, *δ*(*t*) is the Dirac delta-function and *δ*_*ij*_ the Kronecker delta. The parameters *v*_||*σ*_ and *ω*_*σ*_ then represent the magnitude of such stochastic fluctuations, with the translational and rotational diffusion constants are denoted by 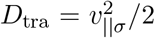 and 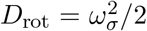 respectively. Finally, the functions *f* (*θ*(*t*), *s*(*t*), *t*) and *g*(*s*(*t*), *t*) are set separately accordingly to the diatom’s motility state. Note that the evolution of the position is coupled to that of the orientation, as *f* (*θ*(*t*), *s*(*t*), *t*) is a function of *θ*(*t*). Moreover, both functions depend on the diatom’s ‘polarity’ *s*(*t*) ∈ {−1, +1}, an indicator representing whether the diatom is moving forwards or backwards (this is an arbitrary choice since diatoms have no identifiable ‘front’ or ‘back’) that switches its value with every reversal.

In the Glide state *f* (*θ*(*t*), *s*(*t*), *t*) = *s*(*t*)**u**_||_(*t*)*v*_||*µ*_ and *g*(*s*(*t*), *t*) = *s*(*t*)*ω*_*µ*_, where *v*_||*µ*_ is the mean gliding speed and *ω*_*µ*_ the mean rotational speed. The diatom moves forwards (or backwards, depending on *s*(*t*)) with additive Gaussian white noise, and rotates with similar dynamics. For simplicity, we assume translational motion is only in the tangential direction (aligned with the cell’s long axis), consistent with the data.

In the Pivot state 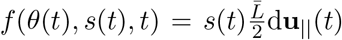, where 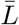 is the mean diatom length of each species, and *g*(*s*(*t*), *t*) = *s*(*t*)*ω*_*µ*_: the orientation changes according to a mean rotational velocity and Gaussian white noise, while the tangential displacement results from rotation about either end of the cell, determined by *s*(*t*).

Lastly, in the Stop the Reverse states *f* (*θ*(*t*), *s*(*t*), *t*) = 0 and *g*(*s*(*t*), *t*) = 0, leaving only the stochastic terms with zero expected net displacement/rotation. The polarity *s*(*t*) switches sign at the transition to Reverse, forcing the subsequent Glide to occur in the opposite direction. Here Reverse is instantaneous, which is an approximation of the actual reversal dynamics.

The values of *v*_||*µ*_, *v*_||*σ*_, *ω*_*µ*_ and *ω*_*σ*_ depend on the the motility state as well as the species, and must be calibrated to the experimental data. We further assume, denoting the associated state by superscripts, 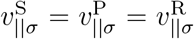 and 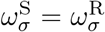, *i*.*e*. translational fluctuations have the same magnitude in Stop, Pivot and Reverse, while rotational fluctuations have the same magnitude in Stop and Reverse. We are then left with a total of 8 parameters to calibrate for each species: 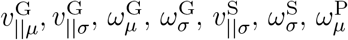, and 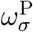.

### Model calibration

We calibrated the model separately on each species. For Glide, we estimated *v*_||_(*t*) and *ω*(*t*) from all tracks, discarding frames not classified as Glide. Since each sequence of consecutive Glide frames has a slightly different mean speed across the different tracks, we perform a velocity scaling on *v*_||_(*t*) to better capture velocity fluctuations *during* glides (see Methods). The same scaling is applied to *ω*(*t*).

Since diatoms have no discernible ‘front’ or ‘back’, reversals lead to a double-peaked distribution for *v*_||_(*t*). Therefore, a generic model should not be biased towards a specific tangential direction (forwards vs backwards) nor a specific rotational direction (clockwise vs counterclockwise), so we count both ±*v*_||_(*t*) and ±*ω*(*t*) to construct distributions that are symmetric about the y-axis (e.g. Fig. 5a for *Seminavis*, results for other species can be found in SI Fig. 8).

**Figure 5.**
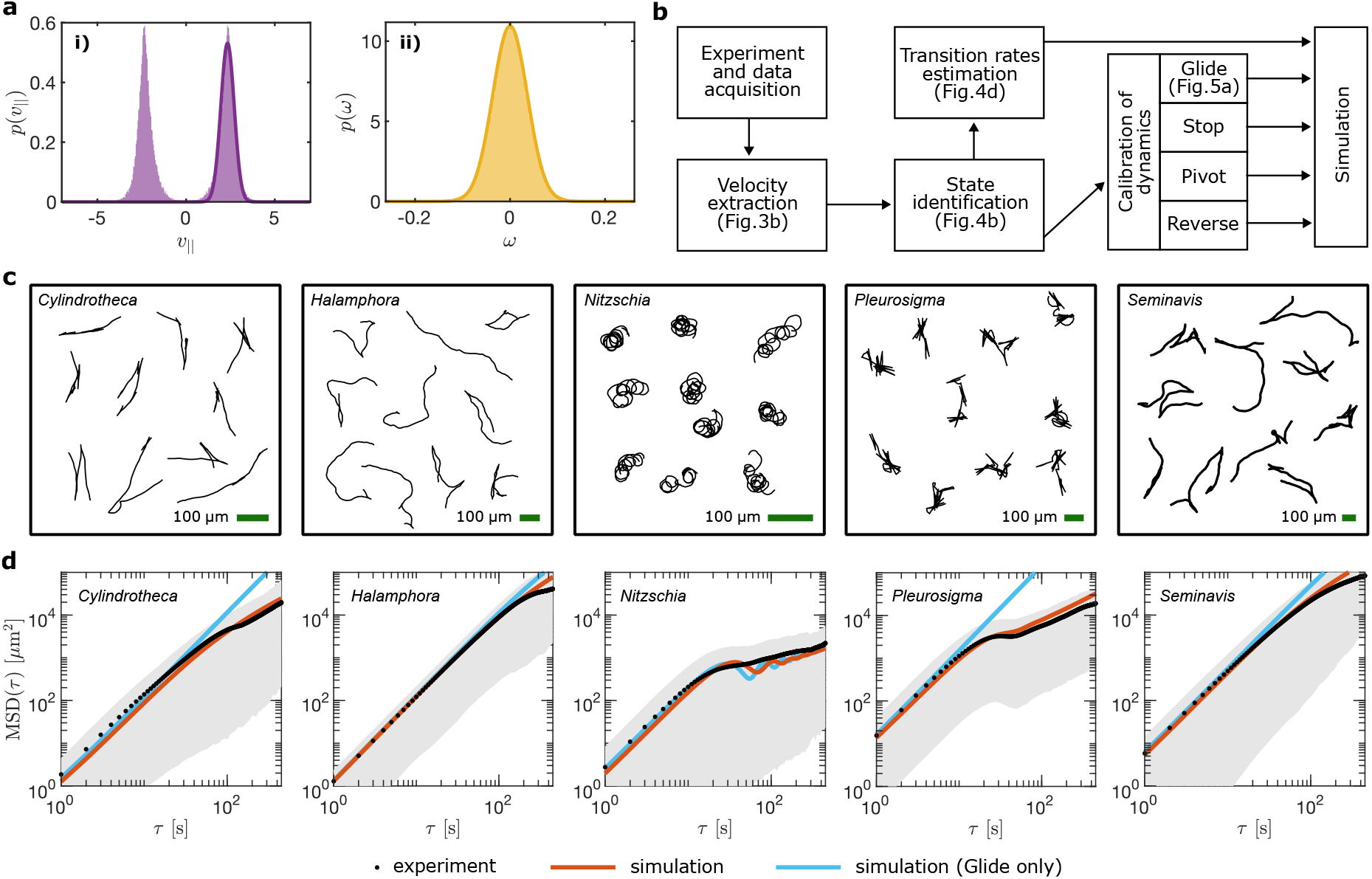
Model calibration and simulation. **a** Calibration of Glide parameters for diatom species *Seminavis*. i) the sum of two normal distributions is fitted to the probability distribution of *v*_||_ to estimate 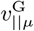 and 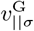. ii) a normal distribution is fitted to the probability distribution of *ω* to estimate 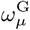 and 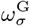. **b** Summary of workflow from experiment to simulation. **c** Examples of simulated trajectories trajectories. **d** MSD obtained from simulations (red line) plotted on top of the experimental MSD (black), with the grey area around it representing the 10-to-90 percentile range of the square displacement. Additional simulations with Glide as the only possible state (cyan).

We improved the fit accuracy with a coarse-graining approach to reduce the influence of measurement noise, using a moving window of 5 frames (see Methods). 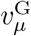 and 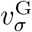 are then fitted to the sum of two Normal distributions 𝒩 (*µ, σ*) and 𝒩 (−*µ, σ*) (Fig. 5a.i):

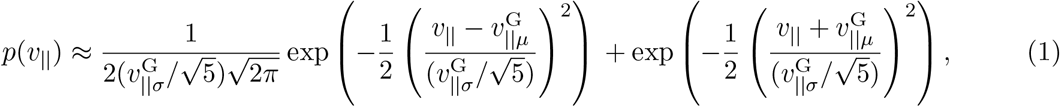

where the factor 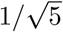 rises from the coarse-graining. For all diatom types except *Nitzschia*, 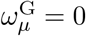, and 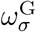 is estimated by fitting to 𝒩 (0, *σ*) (Fig. 4a.ii)

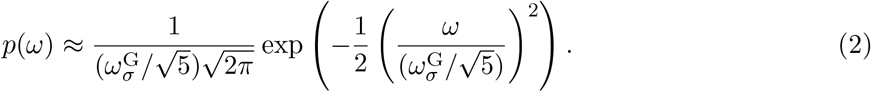

Since *Nitzschia* exhibits chiral motion, 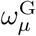 and 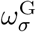 are fitted to the sum of two Normal distributions, analogously to the calibration of 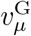 and 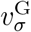. The Stop and Pivot parameters are calibrated analogously (see Methods).

The complete procedure is summarised in Fig. 5b, with distributions of *v*_||_ and *ω* for all diatom species and states, provided in SI Figs. 8-10.

### Long-time dynamics and dispersal

Next, we investigate how the microscale strategy affects macroscale dispersal, in both experiments and simulations. Diatom movement was simulated using a hybrid Gillespie-Langevin algorithm that captures both the discrete-time stochastic switching of the motility state and the continuous-time dynamics of the state itself. The simulated and experimental trajectories qualitatively match (Fig. 5c and Fig. 2), with good quantitative agreement between the Mean Squared Displacement MSD(*τ*) = ⟨∥ **r**(*t*_*i*_ + *τ*) − **r**(*t*_*i*_) ∥_2_⟩ (Fig. 5d).

The MSD analysis shows that all diatoms are ballistic at short times, but diffusive at longer times. *Cylindrotheca* transitions to diffusive motion after ~ 100 s, while textitHalamphora and *Seminavis* transition after a longer time, with observations limited by the maximum recording duration (600 s). To understand whether the ballistic to diffusive transition is due to rotational fluctuations or motility state switching, we performed simulations which do not allow diatoms to leave the Glide state (Fig. 5d). The resulting MSD curves are approximately ballistic up to timescales that are much longer than the maximum duration of the experimental trajectories, showing that the transition to diffusive behaviour can be attributed to reversals and pivots. Since glides are exponentially distributed in *Cylindrotheca, Halamphora, and Seminavis* (Fig. 4e), reversals cause these diatoms to perform a 1D random walk. In contrast, *Pleurosigma* and *Nitzschia* do not transition from ballistic to diffusive behaviour directly. Rather, these species exhibit an intermediate timescale at which the MSD grows sub-linearly due to oscillations: loops in the case of *Nitzschia* and periodic reversals in the case of *Pleurosigma* (Fig. 4e). The period of both loops (~ 41 s, see SI Fig. 11) and reversals (~ 23 s, see Fig. 4e) is shorter than the timescale for randomisation of direction by rotational diffusion, causing these diatoms to revisit recently explored areas. At even longer timescales, diffusive behaviour is recovered (Fig. 6a). Note the discrepancy observed in some diatoms between experimental and simulated MSDs at the highest values of *τ* is due to experimental tracks with large displacement being interrupted when a cell leaves the field of view; the surviving tracks are therefore biased towards smaller displacements. Alternative MSD plots are provided in SI Fig. 12 where only tracks that reached the full 600 seconds are considered.

**Figure 6.**
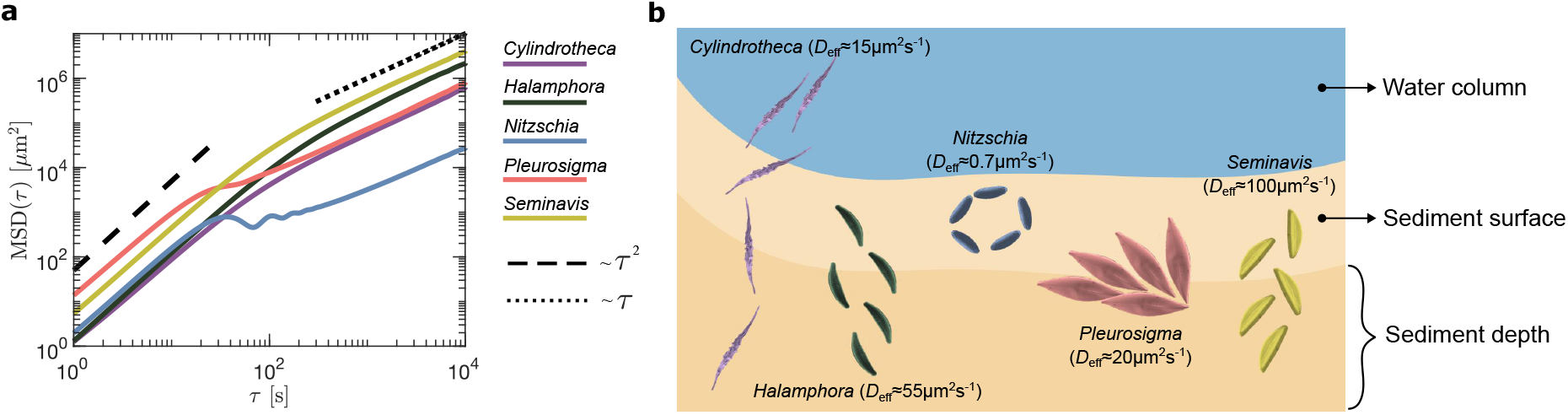
Long-time dispersal and vertical migration. **a** MSDs of all species over timescales of up to 10^4^ s, predicted from the fully-calibrated simulations. The velocity autocorrelation function (see Methods) for the same simulations is shown in Supplementary Fig. 13. **b** Schematic of diatoms’ vertical migration highlighting the different ecological niches they inhabit.

Finally, we predict the overall migration trend for each species with long-time simulations for *τ* up to 10^4^ s (Fig. 6a). Taken together, the results reveal how the distinct motility patterns of the diatoms lead to a natural segregation of their macroscopic diffusivities. These are 15.16 µm^2^ s^−1^ for *Cylindrotheca*, 54.62 µm^2^ s^−1^ for *Halamphora*, 0.70 µm^2^ s^−1^ for *Nitzschia*, 19.64 µm^2^ s^−1^ for *Pleurosigma*, and 100.06 µm^2^ s^−1^ for *Seminavis*, with *Seminavis* and *Halamphora* being the most diffusive, and *Nitzschia* the least diffusive. These are consistent with the putative dispersal of these species from available literature (Fig. 6b; Discussion).

### 3D experiments and simulation

To bridge our findings on controlled 2D substrates with the more naturalistic settings inhabited by the organisms, we created a structured 3D environment (depth ~ 2 mm) using crushed cryolite, a material with a similar refractive index as seawater, enabling direct microscopic observation and 3D tracking (Fig. 7a, and Methods). In this novel assay, cells navigated the interstices and across the surfaces of the cryolite particles, thereby recreating their movement patterns within natural 3D biofilms.

**Figure 7.**
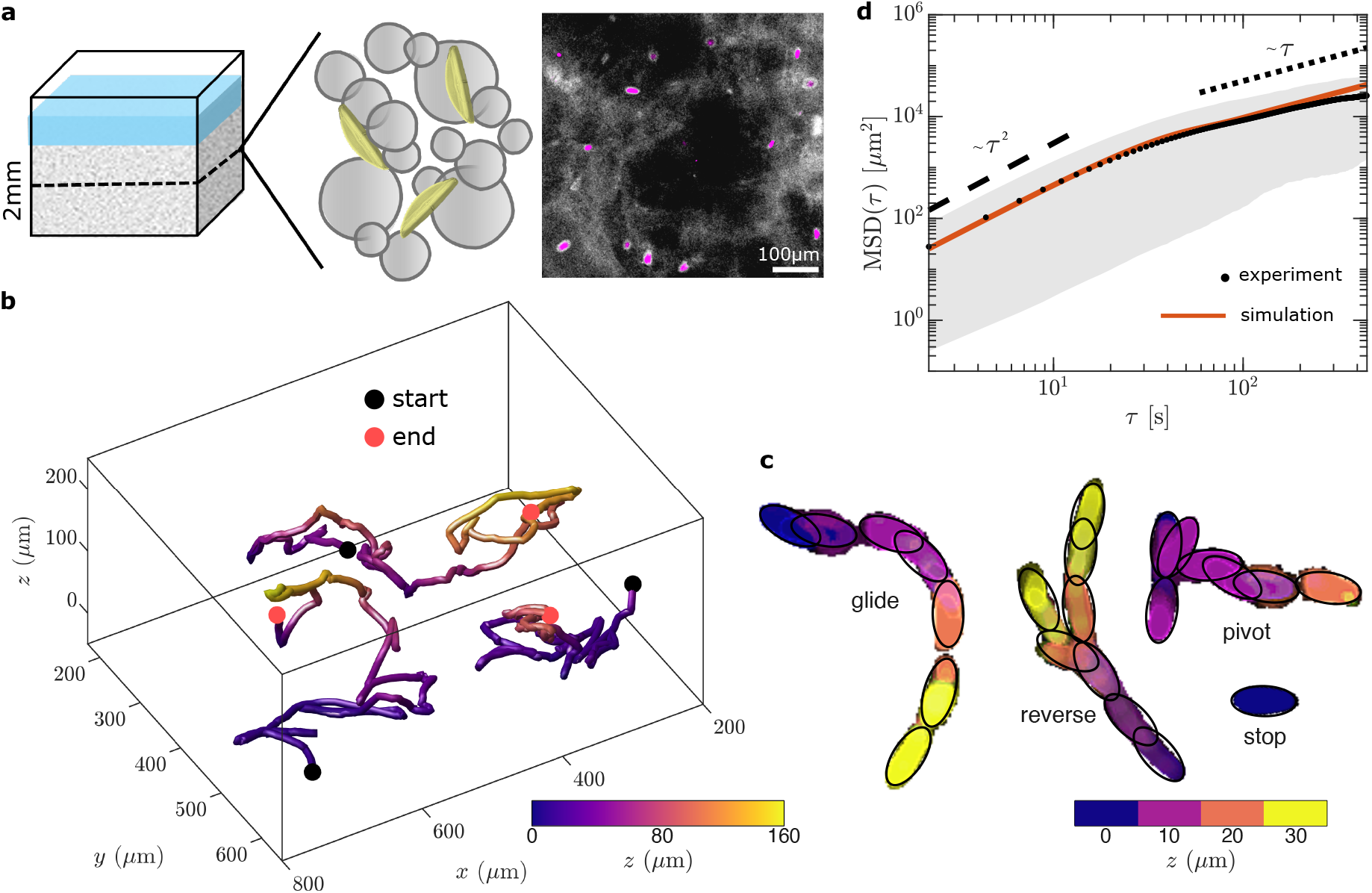
Diatom gliding motility in a complex 3D environment. **a** 3D environment design. Schematic of a 3D substrate (total depth: 2000–2500 *µ*m) made up of cryolite (grey circles) which has a refractive index similar to seawater. The observation area was limited to 200 *µ*m depth. Diatoms can occupy the interstices between and glide along the surfaces of the cryolite particles. Fluorescence image showing the distribution of cells (magenta) in the 3D environment (grey background). Scale bar is 100 *µ*m. **b** Sample tracks of *Seminavis* gliding motility in 3D. **c** *Seminavis* exhibits the four behavioural states in 3D, each colour represents a separate 10*µ*m z-slice. **d** Mean-squared displacement curves from the 3D experimental data and simulations.

We focus on *Seminavis*, chosen because it exhibited all four behavioural states with reasonable probability in our 2D experiments (Fig. 4d), and its larger size facilitates observation. Cells traversed the structure, migrating over vertical distances of ~ 200 *µ*m. Sample 3D tracks are shown in Fig. 7b. As in the 2D experiments, cells showed all four behavioural states of glide, reverse, pivot, and stop (Fig. 7c, SI Video 6). Cells also used pivots and reversals to reorient themselves.

To quantify diatom dispersal in 3D, we again measured the MSD in the experiments (Fig. 7d) (*n* = 151). For *Seminavis*, the 3D MSD matched the 2D curve (Fig. 5d) at small times, but the cells transitioned more rapidly from ballistic to diffusive behaviour in 3D, likely due to collisions with the cryolite particles. To verify this, we extend our simulations to 3D (see Methods), using the same state transition parameters as before, and a new mean glide speed of ~ 2.7*µ*m s^−1^ estimated from the 3D trajectories. To mimic collisions with the obstacles, we stochastically set an upper bound for the dwell time in the Glide state (see Methods). A mean bound time of 37 s was chosen so that diatoms, gliding at an average speed of ~ 2.7*µ*ms^−1^, approximately cover 100*µ*m, the approximate size of the ‘free space’ between particles (*e*.*g*., the white areas in the fluorescence image in Fig. 7a).

These results show that diatom movement through complex 3D environments is quantitatively well-captured by the key features measured in our controlled 2D assays, and suggest that such patterns are intrinsic to each diatom species and therefore a reliable predictor of macroscopic dispersal.

## Discussion

In this contribution, we have studied the gliding motility of five representative biofilm-forming diatoms and revealed their distinctive single-cell motility patterns. Using long-time, high-resolution imaging we discretized their motion into four states based on mathematically well-defined movement characteristics. We developed stochastic simulations to fully capture all features of diatom movement to make predictions about their long-time dispersal. This was extended to 3D, and validated against experimental measurements of cells navigating through realistic 3D environments using a representative diatom. Using emerging analytical and experimental techniques, we have uncovered the natural diversity in behaviour across different motile taxa, and presented a comprehensive new framework that enables precise and quantitative comparisons.

The mysterious gliding motility of diatoms has fascinated researchers for centuries. Gliding has unique features that deviate from established behaviours of flagellated or ciliated microswimmers [5, 39, 41]. The molecular mechanism of diatom motility is hypothesized to involve the raphe and raphe-associated actin bundles [21, 23, 24]. Here, leveraging the distinct raphe morphologies of different diatoms, we have demonstrated that different motility patterns result from the intrinsic shape and curvature profile of the raphe slit. The dominant curvature of gliding trajectories coincides with the curvature of the raphe itself. We therefore expect morphological similarity to translate into similarity in terms of motility phenotype. Indeed, *Halamphora* and *Seminavis* both have simple straight raphes [38] and quantitatively most similar motility patterns. Thus, the functional morphology of the raphe holds essential clues about how motile diatoms diversified throughout evolution, complementing existing taxonomic or phylogenomic evidence.

Our results strongly support the emerging view that motility relies upon an actin-myosin ‘highway’ that is closely aligned with the raphe slit geometry [21, 23, 22]. This mechanism is distinct from those adopted by other gliding eukaryotic cells. In the diatom *Craspedostauros australis*, bidirectional movement of fluorescently-tagged myosins correlates with body movement [24]. Here by simultaneously tracking cell position and body orientation, we showed that cells glide persistently only when the body is aligned with the trajectory heading. Misalignment is associated with reversals or pivots, likely due to changes in adhesion [24, 15]. Out-of-plane tilting movements are most pronounced in diatoms with more complex raphes that cannot adhere fully to the substrate, such as *Cylindrotheca* and *Nitszchia* [42].

Within diatoms, evolution of the raphe and gliding motility was highly consequential for species diversification [43]. Although the raphe places fundamental physical constraints on motility that presumably accounts for the four core behavioural states, we still measured very different diffusivities across species. This functional divergence suggests a possible link between diatom motility, morphology, and evolutionary adaptation to distinct ecological niches. Different motility patterns could lead to niche segregation, influencing the diversification and spatial distribution of organisms [44, 45, 46, 47, 48]. Within mixed-species biofilms, different diatoms, both epipelic — motile cells that live primarily on muddy sediments, and epipsammic — non-motile cells that live among sand grains, are known to occupy specific niches depending on their size, morphology and growth preferences (Fig. 6b). Smaller epipelic diatoms like *Halamphora* or *Seminavis* exploit fast motility for diel migration, traversing the entire photic zone in 0.5 − 1 hour. Larger cells like *Pleurosigma* may inhabit the subsurface where they do not traverse large distances but instead exploit their large body size by constantly reorienting or pivoting their body axis to regulate light exposure [49]. *Nitzschia ovalis* is likely an epipsammic diatom found on the biofilm surface [50, 51] that contributes to overall biofilm cohesion and integrity, where again long-distance migration is not required, consistent with their low diffusivity.

We also obtained the first measurements and direct visualisation of vertical movement in diatoms, revealing that the intrinsic behaviours of gliding diatoms in homogeneous, controlled 2D environments translate readily into complex 3D settings. In nature, distinct motility strategies enable cells to efficiently exploit critical resources, like light, while reducing competition from conspecifics [52, 8, 53]. In the photic zone, which measures ~500 *µ*m in muddy or ~3mm in sandy sediments [54, 55, 28], diatoms use both in-plane and vertical motility to reap the benefits of light for photosynthesis while avoiding light stress [52, 8, 49, 53, 56]. Motility is also influenced by temperature, with ice-dwelling diatoms exhibiting specialised motility strategies to navigate extreme temperatures [15]. More generally, physical constraints can also drive significant phenotypic diversity in other motile organisms [57, 58].

Together, our experiments, analysis and simulations reveal how variability in motility strategy directly influences the macroscale dispersal of diatoms. Using experimentally-calibrated simulations we can confidently predict long-term trends in regimes that are inaccessible in experiments. Future work should explore how these trends may be influenced by environmental perturbations such as light, fluid flow, chemicals, pheromones [11, 8, 53]. Ultimately, our findings should be framed against the scale and environment in which diatoms live to reveal the ecological significance of their motility.

## Methods

### Diatom culturing and experimental set-up

We selected five representative diatom taxa with different frustule shapes and raphe systems that might impact adhesion and motility (Fig. 2A). The diatom strains used are *Cylindrotheca closterium* (DCG 0980), *Halamphora sp*. (CCAP 1050/13), *Nitzschia ovalis* (CCAP 1052/12), *Pleurosigma sp*. (RCC 6814) and *Seminavis robusta* (DCG0096-0115). All diatoms were cultured in f/2 media supplemented with silicate [59] under a 14:10 light:dark cycle at 21°C, with the exception of *Pleurosigma*, which was cultivated at 15°C.

*For 2D experiments*, cultures were prepared by aliquoting stock cultures to an initial cell density of 10^3^ using fresh culture media (total volume per well: 1.5 mL) in *µ*-Slide 2 Well chambers (Ibidi GmbH). The biofilm is considered fully developed after 7-9 days (*>*5×10^4^ cells mL^−1^), videos were recorded to capture baseline motility patterns for that species, in the absence of any other environmental perturbations. We used an inverted microscope (Leica DMi8) equipped with a Leica camera (DMC2900) to take 10 min videos at 1 fps of different observation regions in the well. The total number of videos for each diatom were: *Cylindrotheca* (n=39), *Halamphora* (n=30), *Nitzschia* (n=46), *Pleurosigma* (n=31), and *Seminavis* (n=30).

*For 3D experiments, µ*-Slide 8 Well (Ibidi GmbH) chambers were filled with crushed cryolite (400 mg) to mimic naturalistic 3D environments that are 2000 − 2500 *µ*m in depth. The crystal has a similar refractive index as seawater (*n*_seawater_ = 1.34, *n*_cryolite_ = 1.34), allowing direct microscopic observation of cells in brightfield. To visualise the structure of the 3D environment formed by the powdered crystal (7a), 1 *µ*L of 1:1000 fluorescein (Merck) was added to a well and incubated for 10 mins before observation. Chlorophyll a (blue excitation:450 nm) and fluorescein (green excitation: 550 nm) were consecutively imaged to differentiate between the environment and diatoms.

Live-imaging was performed using *Seminavis* as a representative species. As before, cells were cultured in flasks (TC flasks, 25 cm^2^, Corning) for 7 days reaching a density of *>*5×10^4^ cells mL^−1^. For motility assays, 300 *µ*L of this culture was transferred to each *µ*-Slide 8 Well. Cells were allowed to settle into the wells for a day prior to imaging. To reconstruct the 3D movement of diatoms, we imaged a ~ 200 *µ*m-thick region in the centre of the substrate, obtaining a high-resolution z-stack at 2.2 fps (with 20 slices, stepsize 10 *µ*m) using an inverted microscope (Olympus IX83) coupled to a Kinetix camera (01-Kinetix-M-C, Teledyne).

### Scanning electron microscopy and raphe tracing

For SEM, the samples were initially fixed in a solution comprising 2% glutaraldehyde and 2% paraformaldehyde in 0.1M PIPES buffer at pH 7.2 at room temperature for 1 hour. Following fixation, the cells underwent three washes in the same buffer, each for 5 minutes, and were subsequently transferred onto poly-L-lysine-coated coverslips. Post-fixation was carried out in 1% aqueous osmium tetroxide for 1 hour. This was followed by another series of three 5-minute washes in deionized water. The cells were then subjected to a graded ethanol dehydration series, involving sequential immersions in 30%, 50%, 70%, 80%, 90%, and 95% ethanol for 5 minutes each, followed by two 10-minute immersions in 100% ethanol. Complete dehydration was achieved using hexamethyldisilazane (HMDS, Merck, Gillingham, UK) for 3 minutes, after which the samples were air-dried. The dried samples were mounted onto aluminium stubs using carbon adhesive tabs and subsequently coated with a 10 nm layer of gold/palladium (80/20) using a Q150TES sputter coater (Quorum Technologies Ltd, Laughton, UK). To enhance conductivity, silver paint was applied to the sides of the coverslips. The specimens were then imaged using a Zeiss GeminiSEM 500 scanning electron microscope (Carl Zeiss Ltd, Cambridge, UK) operated at 1.5 kV.

From SEM images, manual raphe shape tracing was done by creating several point selections (n= 50 − 150) along the raphe using the open-source software Fiji [60]. This give a series of *x* and *y* coordinates for each raphe that were used to calculate the raphe curvature (*κ*_*r*_) (see Methods - curvature calculation), Fig.2a and SI Fig. 2).

### Video processing and tracking

We wrote custom Jython scripts in the open-source software Fiji [60] for pre-processing raw videos and tracking. For 2D experiments, cells were detected and tracked as ellipses using the Kalman tracker algorithm in Trackmate [61, 62], the settings of which were optimised for each video. Only extremely motile cells, those that moved at least 15 times their body length, were considered for the analysis (SI Fig. 1). The total number of tracks for each diatom were: 110 for *Cylindrotheca*, 605 for *Halamphora*, 122 for *Nitszchia*, 450 for *Pleurosigma*, and 834 for *Seminavis*.

For 3D experiments, tracking is more challenging, as cells change shape as they go in and out of focus, and can only be detected in Trackmate as spherical spots and could not be fitted to ellipses to obtain cell orientation [62]. We performed spot-tracking using Kalman tracker with similar settings for all videos, to obtain coordinates for the diatom’s position. The resulting tracks were manually inspected to ensure correctly segmented tracks were connected and spurious connections were deleted. We tracked 3 videos resulting in 151 tracks.

To investigate whether diatoms exhibit the same four behavioural states during 3D exploration, we processed the videos by colour-coding each slice or depth (10 *µ*m). Maximum intensity projections were then performed to condense the data into a 2D representation (see SI Video 6, Fig. 7c).

### Calculation of track parameters

For every recorded trajectory, we obtained the time series of the diatoms’ position, with a time interval of Δ*t* = 1s between consecutive frames. The position consists of the two Cartesian coordinates **r**(*t*) = [*x*(*t*), *y*(*t*)] for the centre of the diatom, as well as the diatom’s orientation *θ*(*t*), relative to the *x*-axis taken from the fitted ellipse (Fig. 3a). Due to the symmetry of the ellipse, two possible (opposite) orientations can be identified at each frame. We correct frame-to-frame inconsistencies by assigning *θ*(*t*) ⇒ (*θ*(*t*) + *π*) mod 2*π* when |*θ*(*t*) − *θ*(*t* − 1)| *> π/*2 (*i*.*e*., the smallest rotation is assumed to be the most likely).

At each time frame, the translational velocity was extracted as **v**(*t*) = (**r**(*t* +Δ*t*) −**r**(*t*))*/*Δ*t*, and the angular velocity as *ω*(*t*) = (*θ*(*t* + Δ*t*) − *θ*(*t*))*/*Δ*t*. The velocity **v**(*t*) is decomposed into a tangential (*i*.*e*., in the direction of *θ*(*t*)) and a normal (*i*.*e*., orthogonal to *θ*(*t*)) components. Consider the unit vectors **u**_||_((*t*) = [cos(*θ*(*t*)), sin(*θ*(*t*))] and **u**_⊥_(*t*) = [− sin(*θ*(*t*)), cos(*θ*(*t*))], representing the diatom’s tangential and normal directions, respectively. The velocity is then divided (see Fig. 3b) into a tangential contribution

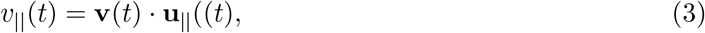

with · the dot product, and a normal contribution

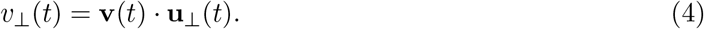

### Curvature calculation

The curvatures *κ* and *κ*_r_ of the diatoms’ tracks and raphes, respectively, are obtained by fitting a circle to three points at different positions along the raphes/tracks, and taking the inverse of the circle’s radius. The raw data was first resampled in order to have the same spatial resolution across raphes and tracks of the same diatom species. Then, curvature values were calculated running a moving window over the raphe/track, with the three points for the circle fitting being the start, the middle and the end of the window. The spatial resolution, which in turn determines the size of the window, is different for each diatom species, thus producing measurements of curvature that are relative to diatoms’ average body-length. A more detailed description and visualisation of both the resampling and curvature estimation procedures can be found in SI Fig. 4.

### Motility state designation

The automation of the motility state designation requires a preliminary processing of the time sequences of tangential velocity, which is described in SI Fig. 14 and produce an alternative velocity 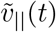. This quantity, alongside *ω*(*t*), is sufficient to reliably automate the state designation via a process that is described in Supplementary Text.

### Estimation of transition probabilities

We implemented a continuous Markov model to describe the cell’s switching between discrete states, designated as Glide, Stop, Pivot and Reverse as outlined above. A transition probability matrix which contains information on when cells move from state *i* to *j* was constructed from the entire dataset. From this, we calculated the pair-wise transition rates between states and the mean dwelling or sojourn time (i.e., the time spent in a state) using the R package *msm* [63].

### Additional details on model calibration

To account for the highly variable speed of gliding, we perform a velocity rescaling. *v*_||_(*t*) is rescaled by the mean glide speed for that glide multiplied by the mean glide speed across all recorded tracks for that species (considering only frames classified as Glide). Doing so, all glides acquire the same mean speed, but the original ratio of their mean to the standard deviation of *v*_||_(*t*) is preserved.

We also perform a coarse-graining approach to reduce measurement noise. Velocities *v*_||_(*t*) and *ω*(*t*) were first randomised to remove temporal correlations and then averaged over a moving window of 5 frames. The dynamics are thus approximated by an effective noise process that ignores all short-timescale variations while producing, on average, equivalent long-timescale random displacements.

For the calibration of 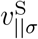 and 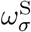, we fit to a Normal distribution with zero mean, as there is no mean displacement/rotation in the Stop state. Instead, 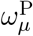 and 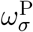 are estimated by fitting the distribution to the sum of two Normal distributions with the same standard deviation and means of equal magnitude and opposite sign, as both clockwise and counterclockwise pivots can occur.

### Model extension to 3D

In the 3D model, the state of the diatom is represented by the position **r**(*t*) = [*x*(*t*), *y*(*t*), *z*(*t*)], the moving frame *E*(*t*) = [*E*_yaw_(*t*), *E*_pitch_(*t*), *E*_roll_(*t*)], where *E*_*i*_(*t*) are orthogonal bases, and the polarity *s*(*t*).

In the Glide motility state, the orientation of the diatom evolves according to

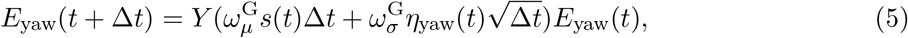

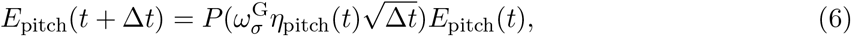

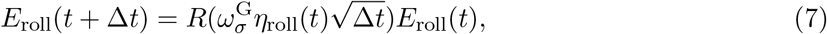

where *η*_*i*_(*t*) are independent sources of Gaussian white noise and *Y* (*α*), *P* (*β*) and *R*(*γ*) are the rotation matrices for the diatom’s yaw, pitch and roll, respectively. Yaw, pitch and roll all involve diffusion, with the assumption that the magnitude of the fluctuations, 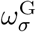, is the same for all rotations. Additionally, yaw involves chiral motion, which we assume only occurs around the normal axis (the axis running from the top to the bottom of the diatom). The position is then updated:

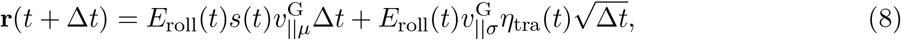

which includes self-propulsion in the direction faced by the diatom, given by *E*_roll_(*t*)*s*(*t*), as well as diffusion along the same axis (*η*_tra_ being the source of Gaussian white noise for translation).

As for the 2D case, Stop and Reverse only involve diffusion, with *s*(*t*) switching sign at the start of a reversal. Pivoting occurs on the diatom’s trailing end and is assumed to be a combination of yaw and pitch. Like the other states, it also included a diffusion component.

To mimic the presence of obstacles in the 3D environment, we set an upper bound on the dwell time in the simulations. After drawing the dwell time from an exponential distribution, another time is also drawn from a Normal distribution with mean 37 seconds (and standard deviation 7.4 seconds), and the final dwelling time is set as the minimum of the two.

## Acknowledgments

This work was funded by UK Research and Innovation (UKRI) under the UK government’s Horizon Europe funding guarantee [grant number EP/X02119X/1] (K.G.B.N and K.Y.W), and the European Research Council (ERC) under the European Union’s Horizon 2020 research and innovation programme grant 853560 EvoMotion (K.Y.W).

We thank Nicole Poulsen, Stefan Diez, Stefan Golfier (Technical University of Dresden) for discussions, Elizabeth Williams (University of Exeter), Georg Pohnert and Franziska Klapper (University of Jena) for providing diatom cultures, Christian Hacker of the Bioimaging Facility for the SEM images of diatoms, and Timothy Naumovitz for assistance with Python codes.

## Author contributions

K.G.B.N. and K.Y.W. designed the research. K.G.B.N. performed the experiments, K.G.B.N., E.C and K.Y.W. analysed the data, E.C. performed the simulations, K.Y.W. supervised the project. All authors discussed the results and contributed to writing and editing.

