## Supplementary Information for "Diversity of motility patterns in benthic diatoms"

### Diversity of motility patterns in benthic diatoms
#### Supplementary Information (SI)

Karen Grace Bondoc-Naumovitz[†,*], Emanuele Crosato[†], and Kirsty Y. Wan[*]

[1]Living Systems Institute & Department of Mathematics and Statistics, Stocker Road, University of Exeter, United Kingdom, EX4 4QD

[†]These authors contributed equally.

#### Supplementary Videos

**Video 1.** Sample video of *Cylindrotheca closterium* cell motility. Individual cells exhibit a glide-reverse movement. The movie was accelerated 50x.

**Video 2.** Sample video of *Halamphora sp.* cell motility. Individual cells exhibit continuous gliding coupled with occasional reverses followed by reorientations. The movie was accelerated 50x.

**Video 3.** Sample video of *Nitzschia ovalis* cell motility. Individual cells exhibit continuous gliding and often form circular tracks. The movie was accelerated 50x.

**Video 4.** Sample video of *Pleurosigma sp.* cell motility. Individual cells exhibit short gliding periods with frequent pivoting behaviours. The movie was accelerated 50x.

**Video 5.** Sample video of *Seminavis robusta* cell motility. Individual cells exhibit continuous gliding coupled with occasional reverses followed by reorientations. The movie was accelerated 50x.

**Video 6.** Sample video of *Seminavis robusta* (SR) cell motility in an artificial 3D environment. Z-stacks were taken at 2.2fps (20 slices, 10$\mu$m apart). Individual cells exhibit continuous gliding coupled with occasional reverses followed by reorientations. The movie was accelerated 50x. Close-up video was accelerated 20x.

#### Supplementary Figures

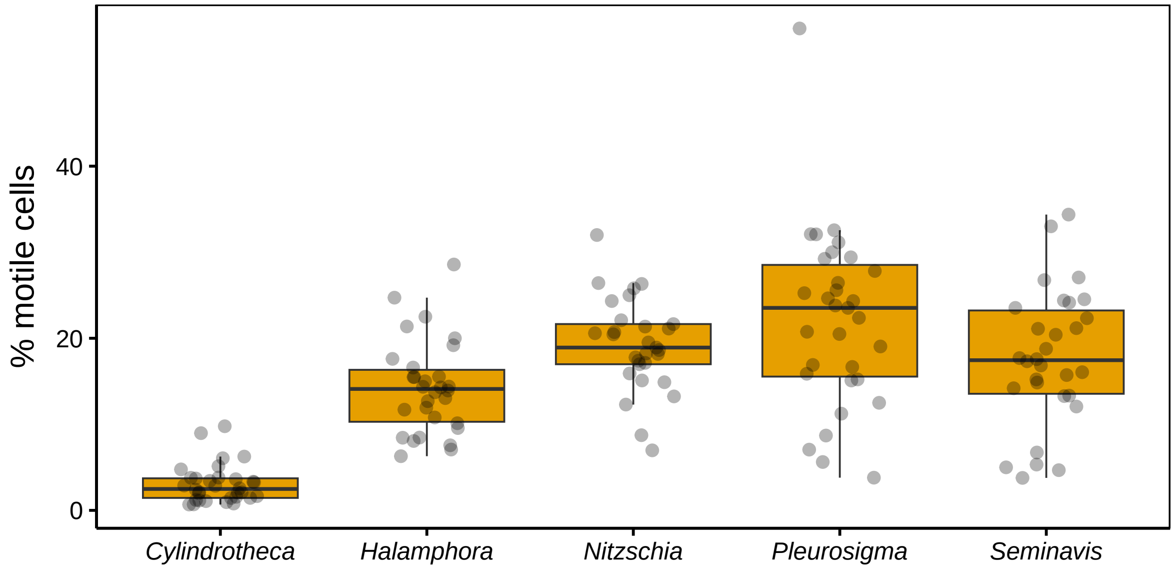


SI Figure 1: **Percentage of motile cells.** Cells that travel 15× more than their body length were used for analysis of 2D motility patterns. Points represent individual videos.

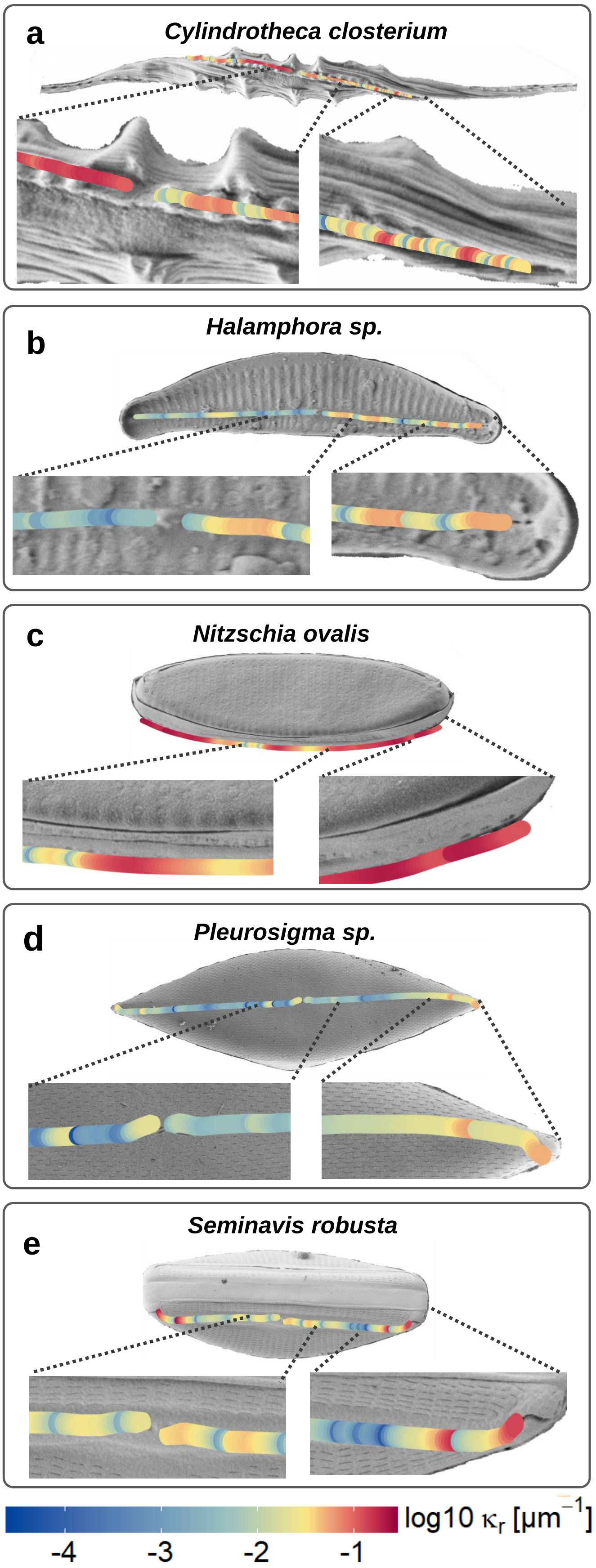


SI Figure 2: **Sample Scanning electron microscopy (SEM) images of the diatoms.** Raphes are traced and coloured by $\kappa_r$ — the measured raphe curvature. Specific regions are highlighted for ease of visualisation. (Also refer to Fig. 2 of the main text.)

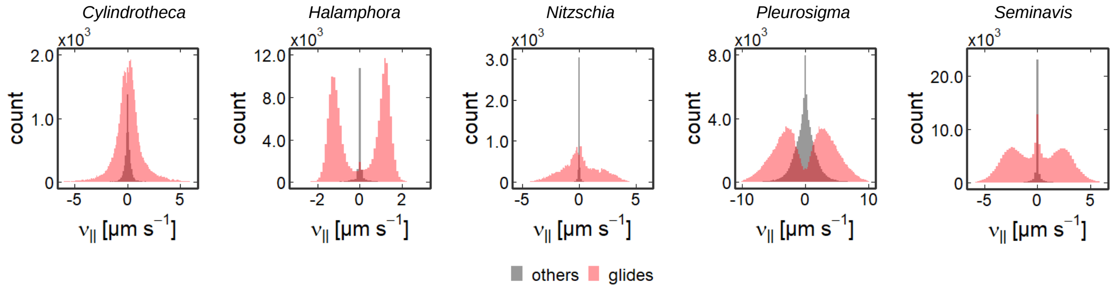


SI Figure 3: **Distributions of $v_{\|}$.** Colours indicate associated $\kappa$ values corresponding to glide and other (*i.e.*, non-glide) movements. Glides were defined based on a specific $\kappa$ threshold for each diatom: *Cylindrotheca* (-0.37 to 0.37), *Halamphora* and *Seminavis* (-0.2 to 0.2), *Nitzschia* (-0.36 to 0.36), *Pleurosigma* (-0.025 to 0.025). For each species, values within the indicated range are considered glides.

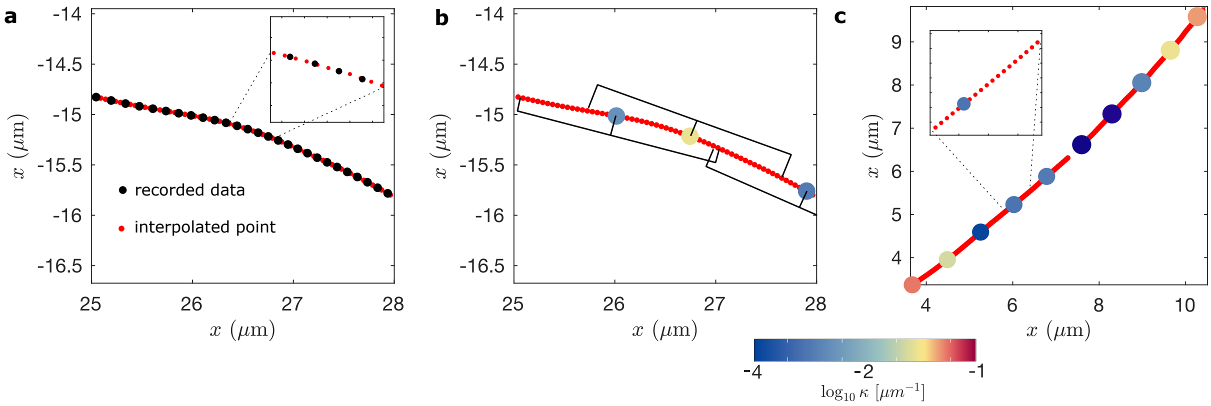


SI Figure 4: **Details of the curvature estimation.** In view of using the same window length for the estimation of the curvature of raphes and tracks, these last need to be resampled at the same spatial resolution. In our 2D experiment, diatoms are recorded at 1 fps, and since their velocity is not constant this results in a sequence of positions that are not equally spaced (black dots, panel (a)). We choose a specific distance of $\bar{L}/300$, with $\bar{L}$ the average diatom length per species. Then, we select the the first recorded position as the current point and repeat the following steps: 1) find the points in the linear interpolation of the track that are at the specific distance from the current point; 2) choose among these the one that is chronologically next and set it as the new current point; 3) repeat the first two steps until the are not more points left. Subsequently, curvature values were calculated by moving a 40-points window over the resampled track, with the three points for the circle fitting being the start, the middle and the end of the window. Three examples measurements are shown in panel (b). Lastly, we repeat the process for the raphes using the same resolution and treating both sides separated by the central nodule individually, as shown in panel (c). Note that the length of the window (which must be the same for raphes and tracks for allowing comparison) has a practical lower bound given by the tracks being recorded at 1 fps: choosing a window length that is smaller than the distance a diatom can travel during a single recording interval would produce invalid measurements. The choice of different window lengths for different species allows a good resolution for smaller diatoms such as *Halamphora* and *Nitzschia*. An exception is made for *Cylindrotheca*, whose complex helical raphe requires a smaller window length and for which a sampling resolution of $\bar{L}/900$ is used instead. This was possible because, despite *Cylindrotheca*'s relatively large size, its average speed is low (see again Fig. 3c and SI Table 1).

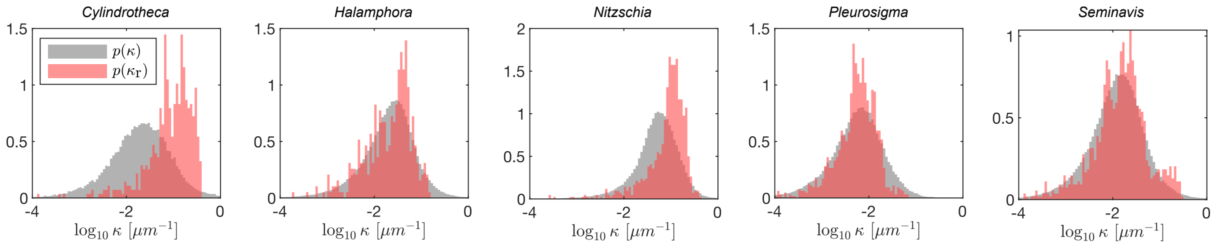


SI Figure 5: **Distributions of glide curvature using all data.** The distribution of the glide curvature, $\kappa$ (gray), is compared with that of the raphe curvature, $\kappa_r$ (red). As discussed in the main text, diatom size varies considerably within the same species and also across species, leading to a broad distribution of curvatures (which we cannot observe in our limited SEM tracings). Note that *Nitzschia* is the most affected by this variance in size/speed.

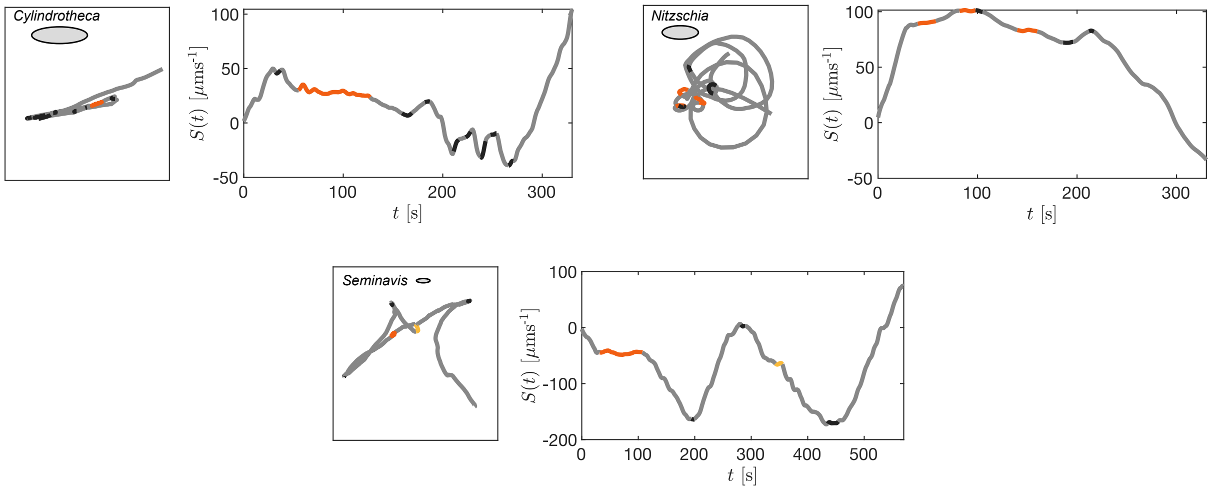


SI Figure 6: **Trajectory segmentation into distinct motility states for *Cylindrotheca, Nitzschia* and *Seminavis*.** The left panels show the trajectories, colour-coded based on the motility state (the ellipses show the average length of the diatoms), while the right panels show the respective time evolution of $S(t)$. (Also refer to Fig. 4b of the main text.)

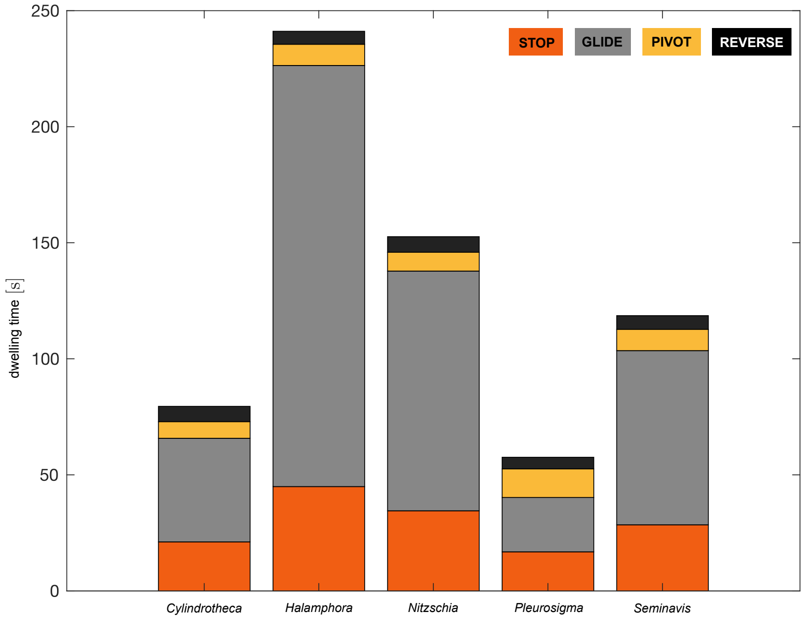


SI Figure 7: **Dwelling times in the four states**. The histograms represent the dwelling time in all states, for all diatom species.)

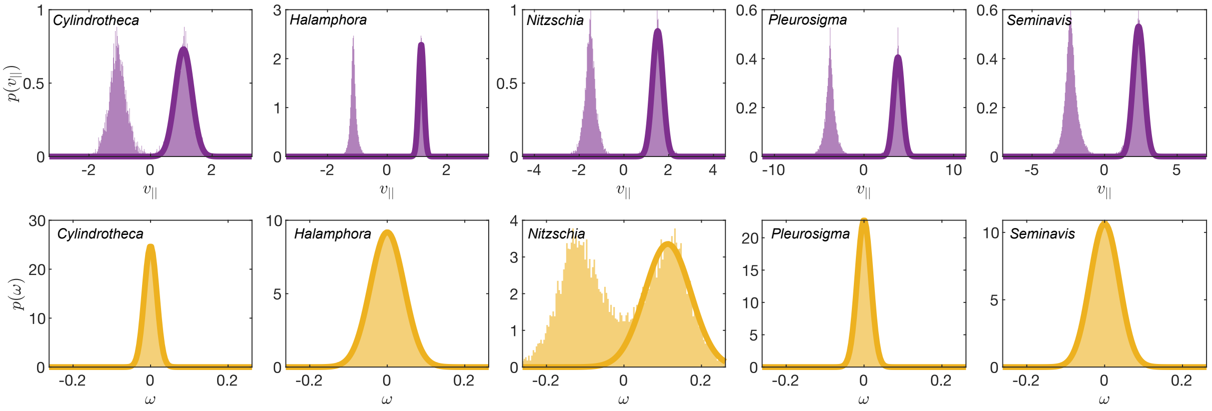


SI Figure 8: **Calibration of Glide parameters.** Top row: fitting of the probability distribution of $v_{||}$ for the calibration of parameters $v_{||\mu}^{\mathrm{G}}$ and $v_{||\sigma}^{\mathrm{G}}$. Bottom: fitting of the probability distribution of $\omega$ for the calibration of parameters $\omega_{\mu}^{\mathrm{G}}$ and $\omega_{\sigma}^{\mathrm{G}}$.

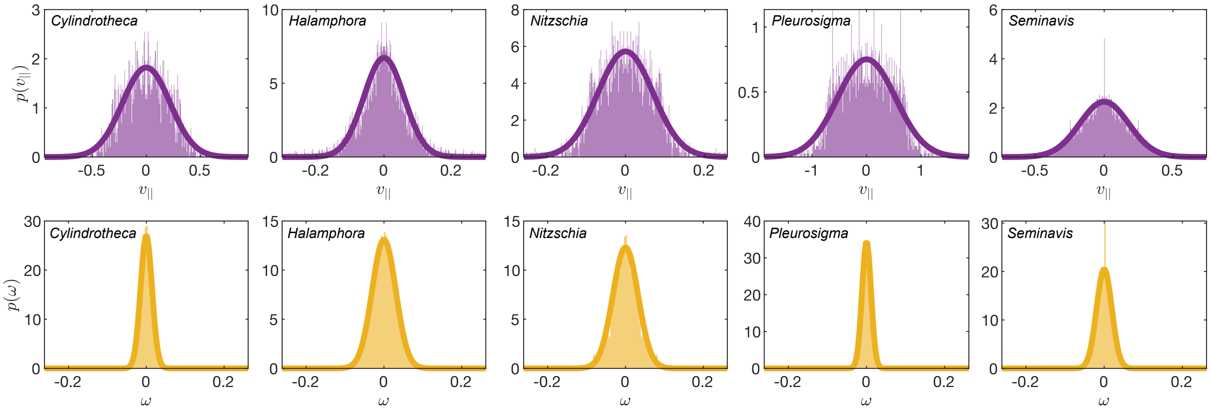


SI Figure 9: **Calibration of Stop parameters.** Top row: fitting of the probability distribution of $v_{||}$ for the calibration of parameter $v_{||\sigma}^{\mathrm{S}}$. Bottom: fitting of the probability distribution of $\omega$ for the calibration of parameter $\omega_{\sigma}^{\mathrm{G}}$.

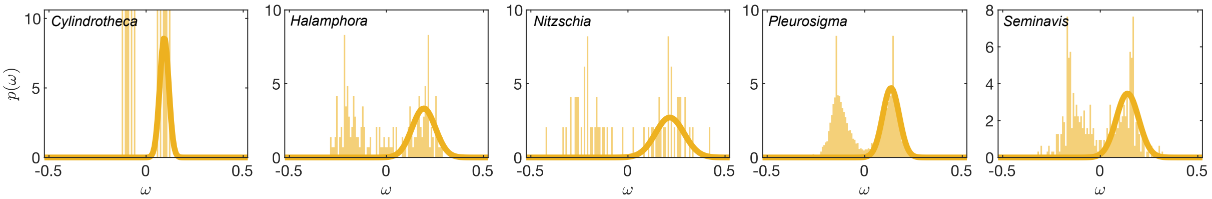


SI Figure 10: **Calibration of Pivot parameters.** Fitting of the probability distribution of $\omega$ for the calibration of parameters $\omega_{\mu}^{\mathrm{P}}$ and $\omega_{\sigma}^{\mathrm{P}}$.

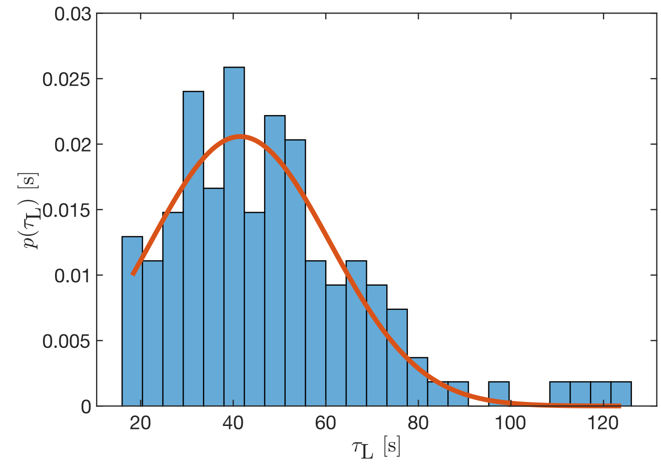


SI Figure 11: **Loop durations for *Nitzschia*.** The histogram shows the distribution of the observed duration of loops $\tau_L$. This is approximated by a truncated Normal distribution with mean 41.52 seconds and standard deviation 19.39 seconds.

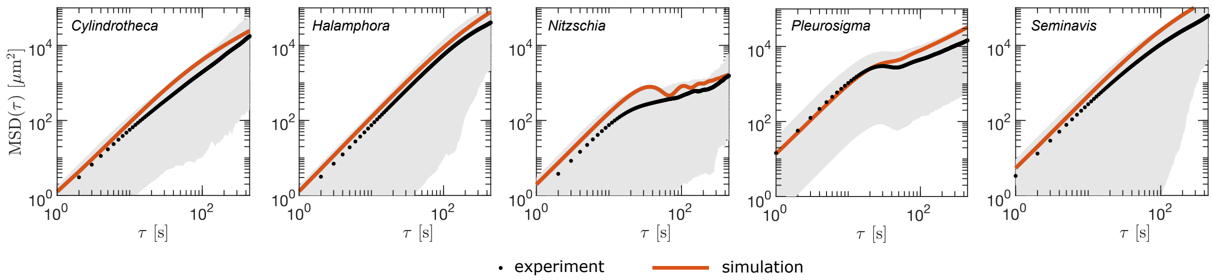


SI Figure 12: **Mean squared displacement from full (600 seconds) tracks only.** The panels show both the experimental and simulated MSD for all diatom species, considering only the tracks that were not interrupted by the cell leaving the experimental field of view. Values of MSD for large $\tau$ are less biased towards slow tracks (those of the diatoms that did not leave the field of view) compared to Fig. 5d in the main text, however, as we are now only considering tracks of smaller displacement corresponding to slower diatoms, the whole experimental MSD line is slightly lower.

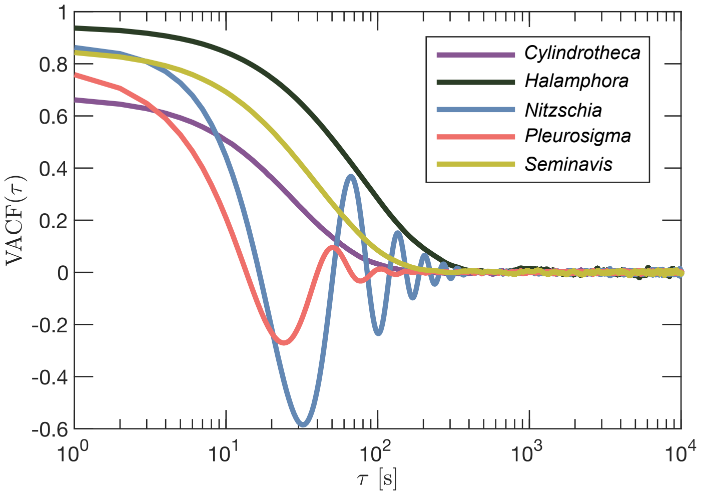


SI Figure 13: **Velocity autocorrelation function.** The VACF is obtained from simulations for all species and shown over timescales of up to 10 000 seconds.

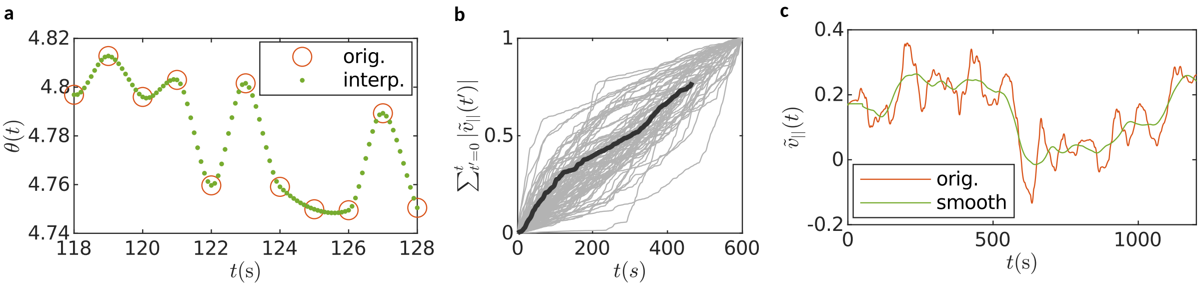


SI Figure 14: **Additional processing steps required for automatic assignment of motility state.** The designation of short events such as reversals and pivots benefits from increasing the sampling of $v_{||}(t)$ from 1 fps to 10 fps. This is done by applying the Makima interpolation separately on $x(t)$, $y(t)$ and $\theta(t)$ and recalculating $v_{||}(t)$ using the interpolated points (panel a). This interpolated velocity is only used for the state designation, any other analysis uses original data exclusively. Since individual diatoms, even within the same species, glide at different preferential speeds, we introduce a *scaled* version of the tangential velocity, defined as

$$\tilde{v}_{||}(t) = \left(\frac{t_{\max}}{T}\right) \frac{v_{||}(t)}{\sum_{t'=0}^{t_{\max}} |v_{||}(t')|},$$

where $t_{\max}$ is duration of a specific track and $T = 600$ seconds is the duration of the longest tracks (panel b). Using $\tilde{v}_{||}(t)$ allows us to use the same classification parameters (*e.g.*, the speed threshold for distinguishing between a glide and a stop). $\tilde{v}_{||}(t)$ is then smoothed as follows: 1) the time series of its cumulative sum $S(t) = \sum_{t'=0}^{t} \tilde{v}_{||}(t')\Delta t$ is obtained, 2) the order one Savitzky-Golay filter with a 10 seconds window is applied to $S(t)$, and 3) the smoothed $S(t)$ is numerically differentiated to obtained a smoothed version of $\tilde{v}_{||}(t)$ (panel c).

### Supplementary Text

#### Algorithm for motility state designation

At every time frame, the average value of the $\tilde{v}_{||}(t)$ is calculated over a time window of 8 seconds centred on the frame. If the value is above a threshold of 0.025/600 then the frame is classified as `Glide`, otherwise it is temporarily classified as `non-Glide`. `Glide` sequences that are shorter than 3 seconds are changed to `non-Glide` and, similarly, `non-Glide` sequences that are shorter than 3 seconds are changed to `Glide`.

The `non-Glide` sequences as then classified as either `Stop` or `Reverse`. If we define $t_1$ as the time just before the start of a `non-Glide` sequence (minus 4 seconds, half of the time widow used for determining the `Glide` states), and $t_2$ as the time right after the end of the `non-Glide` sequence (plus 4 seconds), then $\tilde{v}_{||}(t_1)\tilde{v}_{||}(t_2) \geq 0$ means that the gliding direction was not reversed, and thus the whole `non-Glide` sequence is classified as `Stop`. Otherwise, the duration of the `non-Glide` sequence is inspected: if this is shorter then three times the duration of a typical reversal (which we estimate to be 5 seconds), then the entire `non-Glide` sequence is classified as `Reverse`, otherwise it is classified as a `Stop` sequence followed by a `Reverse` sequence of 5 seconds at the end.

`Pivot` states are identified last. The position of the two ends of the diatom elliptical shape are obtained as $\mathbf{r}_{\pm}(t) = \mathbf{r}(t) \pm (\bar{L}/2)\mathbf{u}_{||}(t)$, and their velocities $\mathbf{v}_{\pm}(t)$ are calculated at each time frame. If $\| \mathbf{v}_{+}(t) \| / \| \mathbf{v}_{-}(t) \| > 2$ or $\| \mathbf{v}_{-}(t) \| / \| \mathbf{v}_{+}(t) \| > 2$, then the time frame $t$ is classified as `Pivot`. Sequences of less than 3 consecutive `Pivot` frames are changed to `Glide`, and sequences of less than 3 consecutive `Glide` frames between `Pivots` are changed to `Pivot`. Every previous classification of states is overwritten by `Pivot` sequences, which are extended to include any `Reverse` occurring right before or after. Then, $\tilde{v}_{||}(t)$ is inspected before and after the extended `Pivot` sequence: if its sign switches, then a `Reverse` sequence is added at the end of the sequence.

### Supplementary Tables

| diatom | cell shape | cell length (mean ±sd µm) | raphe characteristics |
|---|---|---|---|
| *Cylindrotheca closterium* | thin and needle-like | 55.6±9.9 | twisted raphe along its body |
| *Halamphora sp.* | asymmetric biraphid, strongly dorsiventral | 15±1.09 | straight eccentric raphe, positioned along the central margin |
| *Nitzschia ovalis* | ovoid | 8±0.8 | eccentric fibulate raphe lie on opposite sides of the cell |
| *Pleurosigma sp.* | symmetric biraphid, sigmoid-shaped | 75.7±5.2 | centred sigmoid-shaped raphe on each side of the valve |
| *Seminavis robusta* | asymmetric biraphid | 25.6±4.6 | straight raphe displaced laterally |

SI Table 1: Morphology of the five representative diatoms used in this study highlighting the differences between the cell shape and raphe characteristics of each species. Length is taken from n=20 cells for each diatom.

| | STOP | GLIDE | PIVOT | REVERSE |
|---|---|---|---|---|
| STOP | 0.00 | 0.83 | 0.01 | 0.16 |
| GLIDE | 0.37 | 0.00 | 0.01 | 0.63 |
| PIVOT | 0.44 | 0.44 | 0.00 | 0.11 |
| REVERSE | 0.00 | 1.00 | 0.00 | 0.00 |

SI Table 2: Pair-wise transition probabilities of *Cylindrotheca*.

| | STOP | GLIDE | PIVOT | REVERSE |
|---|---|---|---|---|
| STOP | 0.00 | 0.77 | 0.10 | 0.14 |
| GLIDE | 0.17 | 0.00 | 0.07 | 0.77 |
| PIVOT | 0.21 | 0.43 | 0.00 | 0.36 |
| REVERSE | 0.00 | 1.00 | 0.00 | 0.00 |

SI Table 3: Pair-wise transition probabilities of *Halamhora*.

| | STOP | GLIDE | PIVOT | REVERSE |
|---|---|---|---|---|
| STOP | 0.00 | 0.70 | 0.11 | 0.20 |
| GLIDE | 0.34 | 0.00 | 0.06 | 0.60 |
| PIVOT | 0.39 | 0.31 | 0.00 | 0.30 |
| REVERSE | 0.00 | 1.00 | 0.00 | 0.00 |

SI Table 4: Pair-wise transition probabilities of *Nitzschia*.

| | STOP | GLIDE | PIVOT | REVERSE |
|---|---|---|---|---|
| STOP | 0.00 | 0.62 | 0.26 | 0.12 |
| GLIDE | 0.04 | 0.00 | 0.13 | 0.83 |
| PIVOT | 0.17 | 0.19 | 0.00 | 0.64 |
| REVERSE | 0.00 | 1.00 | 0.00 | 0.00 |

SI Table 5: Pair-wise transition probabilities of *Pleurosigma*.

| | STOP | GLIDE | PIVOT | REVERSE |
|---|---|---|---|---|
| STOP | 0.00 | 0.79 | 0.10 | 0.11 |
| GLIDE | 0.27 | 0.00 | 0.05 | 0.68 |
| PIVOT | 0.36 | 0.34 | 0.00 | 0.31 |
| REVERSE | 0.00 | 1.00 | 0.00 | 0.00 |

SI Table 6: Pair-wise transition probabilities of *Seminavis*.

| | mean | s.e. | lower CI | upper CI |
|---|---|---|---|---|
| STOP | 21.11 | 1.18 | 18.92 | 23.56 |
| GLIDE | 44.65 | 1.57 | 41.68 | 47.83 |
| PIVOT | 7.17 | 2.28 | 3.85 | 13.35 |
| REVERSE | 6.58 | 0.28 | 6.06 | 7.15 |

SI Table 7: Mean sojourn or dwell times of *Cylindrotheca*.

| | mean | s.e. | lower CI | upper CI |
|---|---|---|---|---|
| STOP | 44.94 | 2.58 | 40.16 | 50.30 |
| GLIDE | 181.45 | 5.36 | 171.24 | 192.26 |
| PIVOT | 9.15 | 0.90 | 7.54 | 11.10 |
| REVERSE | 5.59 | 0.18 | 5.24 | 5.95 |

SI Table 8: Mean sojourn or dwell times of *Halamphora*.

| | mean | s.e. | lower CI | upper CI |
|---|---|---|---|---|
| STOP | 34.52 | 2.18 | 30.50 | 39.08 |
| GLIDE | 103.29 | 4.42 | 94.97 | 112.34 |
| PIVOT | 8.09 | 1.04 | 6.29 | 10.40 |
| REVERSE | 6.72 | 0.34 | 6.08 | 7.42 |

SI Table 9: Mean sojourn or dwell times of *Nitzschia*.

| | mean | s.e. | lower CI | upper CI |
|---|---|---|---|---|
| STOP | 16.83 | 0.63 | 15.64 | 18.11 |
| GLIDE | 23.41 | 0.25 | 22.92 | 23.91 |
| PIVOT | 12.33 | 0.35 | 11.67 | 13.02 |
| REVERSE | 5.01 | 0.06 | 4.90 | 5.12 |

SI Table 10: Mean sojourn or dwell times of *Pleurosigma*.

| | mean | s.e. | lower CI | upper CI |
|---|---|---|---|---|
| STOP | 28.48 | 0.85 | 26.87 | 30.19 |
| GLIDE | 75.05 | 1.30 | 72.55 | 77.65 |
| PIVOT | 9.15 | 0.53 | 8.16 | 10.25 |
| REVERSE | 5.92 | 0.12 | 5.69 | 6.15 |

SI Table 11: Mean sojourn or dwell times of *Seminavis*.
